# Control of SIV infection in prophylactically vaccinated, ART-naïve macaques is required for the most efficacious CD8 T cell response during treatment with the IL-15 superagonist N-803

**DOI:** 10.1101/2022.06.02.494515

**Authors:** Amy L. Ellis-Connell, Alexis J. Balgeman, Olivia E. Harwood, Ryan V. Moriarty, Jeffrey T. Safrit, Andrea M. Weiler, Thomas C. Friedrich, Shelby L. O’Connor

**Author notes:** ImmunityBio, Culver City, CA 90232. Address correspondence to Shelby L. O’Connor: e-mail. Funding: This study was supported by funding supplied through the National Institutes of Health (NIH R01 grant number AI108415). This research was conducted at a facility constructed with support from Research Facilities Improvement Program grant numbers RR15459-01 and RR020141-01. The Wisconsin National Primate Research Center is also supported by grants P51RR000167 and P51OD011106. Conflict of interest statement: J.T.S. is an employee of ImmunityBio.

## Abstract

The IL-15 superagonist N-803 has been shown to enhance the function of CD8 T cells and NK cells. We previously found that in a subset of vaccinated, ART-naïve, SIV+ rhesus macaques, N-803 treatment led to a rapid but transient decline in plasma viremia that positively correlated with an increase in the frequency of CD8 T cells. Here we tested the hypothesis that prophylactic vaccination was required for N-803 mediated suppression of SIV plasma viremia. We vaccinated rhesus macaques with a DNA prime/Ad5 boost regimen using vectors expressing SIVmac239 gag, with or without a plasmid expressing IL-12, or left them unvaccinated. Animals were then intravenously infected with SIVmac239M. Six months after infection, animals were treated with N-803. We found no differences in control of plasma viremia during N-803 treatment between vaccinated and unvaccinated macaques. Furthermore, the SIV-specific CD8 T cells displayed no differences in frequency or ability to traffic to the lymph nodes. Interestingly, when we divided the SIV+ animals based on plasma viral load set-point prior to N-803 treatment, N-803 increased the frequency of SIV-specific T cells expressing ki-67+ and granzyme B+ in animals with low plasma viremia (<10^4^ copies/mL; SIV controllers) compared to animals with high plasma viremia (>10^4^ copies/mL;SIV non-controllers). In addition, Gag-specific CD8 T cells from the SIV+ controllers had a greater increase in CD107a+CD8+ T cells when compared to SIV+ non-controllers. Overall, our results indicate that N-803 is most effective in SIV+ animals with a pre-existing immunological ability to control SIV replication.

## Introduction

While nearly all individuals mount a CD8 T cell-mediated immune response to acute HIV/SIV infection, very few individuals spontaneously control HIV/SIV replication (1, 2). Several studies comparing the immune responses of elite controllers to those of progressors have implied that polyfunctional CD8 T cells, consisting of both cytolytic and non-cytolytic responses, are responsible, at least in part, for the control of HIV/SIV infection(3–5).

Prophylactic vaccine studies have been performed to elicit polyfunctional CD8 T cells that mimic those present in elite controllers (6, 7). Despite this, most of these strategies fail, and the CD8 T cells lose their potential to kill infected target cells and suppress virus replication (8–10).

Recently, many investigators have turned to the use of immunotherapeutic agents which could, in combination with vaccines, improve the ability of CD8 T cells to target and destroy infected cells during HIV/SIV infection (11). One such class of immunotherapeutic agents that can boost the CD8 T cell response to HIV/SIV is interleukin-15 (IL-15) agonists (12, 13). IL-15 is a cytokine that is normally produced by antigen-presenting cells during viral infections, and promotes the development and growth of several innate and adaptive immune cells, in particular, NK and CD8 T cells(14–16).

One member of the class of IL-15 agonists that has gained enthusiasm in immunotherapeutic studies and clinical trials recently is N-803. N-803 is a soluble IL-15 superagonist, where a constitutively active IL-15 molecule containing a single amino acid mutation (N72D) is bound to the sushi domain of IL-15Rα and fused to the Fc region of IgG1 (17, 18). N-803 is in use in clinical trials in cancer patients, as it improves the ability of CD8 T cells and NK cells to target and destroy tumor cells (19).

There is a growing body of evidence suggesting that N-803 may be an ideal immunotherapeutic agent in HIV+ individuals. Previous *in vivo* studies of SIV+ macaques (20–22) indicated that N-803 treatment increased CD8 T cell and NK cell frequencies. N-803 also increased the frequencies of these cells in the lymph nodes, likely attributed to increased expression of lymph node homing markers such as CXCR5 (21). These promising features of N-803 have been a part of the rationale to test it in a Phase II clinical trial in HIV+ individuals in Thailand under the name of Anktiva™ (https://immunitybio.com/immunitybio-announces-launch-of-phase-2-trial-of-il-15-superagonist-anktiva-with-antiretroviral-therapy-to-inhibit-hiv-reservoirs/), as well as in clinical trials in the United States (23) (https://actgnetwork.org/studies/a5386-n-803-with-or-without-bnabs-for-hiv-1-control-in-participants-living-with-hiv-1-on-suppressive-art/).

While N-803 has the potential to boost the frequency and cytolytic function of CD8 T cells *in vivo* (20–22), the subsequent impact on HIV/SIV replication is less clear. We previously treated four ART-naïve SIV+ rhesus macaques with N-803, and they all exhibited transient control of SIV plasma viremia within 7 days of N-803 treatment (20). Suppression of SIV replication in these four animals, in the absence of ART, did not completely extend to other SIV+ animal studies (21). This could be partly attributed to the latency reversing activity of N-803 (24, 25). Understanding the conditions under which N-803 can successfully boost immune responses to control actively replicating SIV/HIV or reduce the viral reservoir is necessary to improve the clinical relevance of this agent.

Here, we begin to define the features of SIV+ macaques that are associated with N-803-mediated suppression of SIV replication. Previous associations include a lower chronic viral load set point ( ≤10^4^ SIV gag copies/mL plasma) prior to N-803 treatment, host MHC genetics associated with spontaneous SIV control, and prior vaccination (20, 21). In this study, we tested the hypothesis that pre-existing vaccine-elicited CD8 T cells were required for N-803-mediated suppression of SIV replication. We used rhesus macaques (RM) expressing the *Mamu*-A*001 MHC class I allele, which is not associated with natural control of SIV infection (26–28). Unfortunately, treatment with N-803 failed to reduce plasma SIV viremia in vaccinated or unvaccinated macaques.

We then rearranged the animal groups to determine if spontaneous initial control of SIV replication was associated with improved immunological responsiveness of polyfunctional virus-specific CD8+ T cells to N-803. We characterized the function of peripheral SIV-specific CD8 T cells prior to and during the first seven days after N-803 treatment. We found that the proliferative and cytolytic capacity of the SIV-specific cells from the SIV non-controllers did not increase to the same extent as the SIV controllers after N-803 treatment. Our results imply that N-803 efficacy is likely maximized in a host who has not succumbed to severe immunological dysfunction associated with SIV/HIV pathogenesis.

## Materials and Methods

## Animals and reagents

### Animals

All fifteen (15) newly infected Indian rhesus macaques involved in this study were genotyped for the MHC class I allele *Mamu-A1*001* using methods described previously (29). All animals involved in this study were cared for and housed at the Wisconsin National Primate Resource Center (WNPRC), following practices that were approved by the University of Wisconsin Graduate School Institutional Animals Care and Use Committee (IACUC; protocol number G005507). All procedures, such as administration of vaccines, biopsy, and blood draw collection, and N-803 administration were performed as written in the IACUC protocol, under anesthesia to minimize suffering.

Frozen samples were also analyzed from rhesus macaques included in a previously published study (20) that was approved under protocol #G005507. Table I shows a list of the animal ID, sex, and age of each animal, and which figures samples from each animal were included.

**Table I.** Animals included in study.

| Animal ID | Sex | Age (years) | Figures samples were used for |
| --- | --- | --- | --- |
| rh2903 | M | 5.3 | 1-10; S4 |
| rh2906 | F | 4.2 | 1-10; S4 |
| rh2907 | F | 4.4 | 1-10; S4 |
| rh2909 | M | 5.8 | 1-10; S4 |
| rh2911 | M | 5.7 | 1-10; S4 |
| rh2919 | F | 4.8 | 1-10; S4 |
| rh2920 | M | 6.7 | 1-10; S4 |
| rh2921 | F | 4.8 | 1-10; S4 |
| r05053 | F | 15 | 1-10; S4 |
| r15053 | M | 3.8 | 1-10; S4 |
| r15090 | F | 3.6 | 1-10; S4 |
| r03019 | F | 15.3 | 1-10; S4 |
| r10063 | M | 10.3 | 1-10; S4 |
| rh2493 | M | 12.1 | 1-10; S1-S4 |
| rh2498 | M | 12.1 | 1-10; S3,S4 |
| r08016 | M | 8.7 | 5-10; S4 |
| r08054 | M | 8.3 | 5-10; S4 |
| r09089 | M | 7.4 | 5-10; S1, S4 |
| r11021 | M | 5.9 | 5-10; S4 |

S1-S4, Supplementary figures 1-4.

### DNA plasmid vector

The endotoxin-free DNA plasmid vaccine vector (hCMV/R-SIVmac239 *gag*) utilized in this study consisted of the SIVmac239 *gag* gene produced under the control of a CMV promoter (hCMV/R) and was constructed as previously described (30). This plasmid vector was kindly provided by Drs. John Mascola, Robert Seder and Wing-Pui Kong at the NIH Vaccine Research Center.

### Rhesus IL-12 DNA plasmid vector

The endotoxin-free rhesus IL-12 DNA plasmid vector (plasmid AG157) was kindly provided by Drs. George Pavlakis and Barbara Felber. This plasmid vector consisted of the two subunits of the rhesus IL-12 gene expressed from dual promoters in the plasmid and was constructed as previously described (31).

### rAd5 SIVmac239gag vaccine vector

All vaccinated animals received 10^10^ particles of a replication-incompetent rAD5 vaccine vector expressing SIVmac239 gag (ViraQuest; North Liberty, IA).

### N-803 reagent

N-803 was provided by ImmunityBio (San Diego, CA), and produced using methods previously described (18).

## Clinical procedures and viral loads

### Vaccination of Rhesus macaques

The vaccine regimen consisted of a heterologous DNA prime/Ad5 boost strategy. Briefly, all vaccinated macaques received two (2) doses of 2mg of a DNA vaccine vector expressing the SIVmac239 *gag* gene, with or without 0.2mg of a rhesus macaque IL-12 plasmid expression vector, delivered by electroporation (described below). Each dose was separated by four (4) weeks. Four weeks after the second DNA vaccination, animals were given a boost vaccine consisting of 10^10^ particles of an rAD5 vaccine vector expressing SIVmac239 *gag*, delivered intramuscularly.

### Delivery of DNA vectors by electroporation

As described above, animals were given two doses of DNA vaccine vector, delivered approximately 4 weeks apart, following protocols as previously described (32). Briefly, 2mg of hCMV/R-SIVmac239 *gag* plasmid, with or without 0.2mg of rhesus macaque IL-12 plasmid (AG157) were prepared in 500 μL of endotoxin-free PBS. The plasmids were delivered intramuscularly via electroporation. The electroporation device was provided by Ichor Medical Systems (San Diego, CA). The electroporation method has been utilized in previous studies (33) and consisted of an electrical impulse of 40ms over a 400ms duration, at 250V/cm.

### SIVmac239M infection

All fifteen (15) animals in this study were infected intravenously with 10,000 infectious units (IU) of a barcoded SIVmac239, termed SIVmac239M. In the case of the vaccinated macaques, SIV infection occurred eight (8) weeks after the rAd5 vaccination. The SIVmac239M virus used in this study was kindly provided by Dr. Brandon Keele and was constructed as previously described (34).

### N-803 administration

The N-803 dose and route of administration used in this study were previously determined to be safe and efficacious in macaques (18, 20). In the present study, all macaques received N-803 approximately 6 months after SIV infection. They received three (3) doses of 0.1mg/kg N-803, administered subcutaneously. Each dose was separated by two weeks.

### Viral loads

Plasma viral loads were quantified as previously described (20, 35). Briefly, viral RNA (vRNA) was isolated from plasma samples using the Maxwell Viral Total Nucleic Acid Purification kit (Promega, Madison WI). Then, vRNA was reverse transcribed using the TaqMan Fast Virus 1-Step qRT-PCR kit (Invitrogen) and quantified on a LightCycler 480 or LC96 instrument (Roche, Indianapolis, IN).

## IFNγ ELISpot assays

To validate that vaccination elicited Gag-specific immune responses, IFNγ ELISPOT assays were used, as previously described (36, 37). Briefly, frozen peripheral blood mononuclear cells (PBMCs) from the indicated timepoints pre- and post-vaccination were thawed, and 1X10^5^ cells were then added to each well of a monkey IFNγ ELISPOT plate (Mabtech, Sweden), along with 1μM of either Gag_181-189_CM9 or Tat_28-35_SL8 peptides in RPMI media supplemented with 10% Fetal bovine serum (FBS). As negative and positive controls, cells were incubated with either RPMI media with 10% FBS alone, or media supplemented with 10ug/mL concanavalin A, respectively. The plates were incubated overnight at 37°C and 5% CO_2_, and developed the following day according to the manufacturer’s instructions. Plates were read by using an AID robotic ELISPOT reader (AID, Strassberg, Germany). A positive response was defined as the number of spot-forming colonies (SFCs) per 10^6^ PBMCs that were 2 standard deviations above the average value for the negative control, or 50 SFCs/10^6^ PBMC, whichever was greater.

## Flow cytometric analysis

### Tetrameric reagents

The *Mamu*-B*008 Nef_137-146_RL10 and *Mamu*-A1*001 Tat_28-35_SL8 biotinylated monomers were produced by the NIH Tetramer Core Facility at Emory University (Atlanta, GA). The *Mamu*-A1*001 Gag_181-189_CM9 biotinylated monomer was purchased from MBL International (Woburn, MA). The *Mamu*-B*008 Nef_137-146_RL10 and *Mamu*-A*001 Gag_181-189_CM9 monomers were then tetramerized with streptavidin-PE (0.5mg/mL, BD biosciences) at a 4:1 molar ratio of monomer: streptavidin. Briefly, 1/10th volumes of streptavidin-PE were added to the monomer every 10 minutes and incubated in the dark at 4°C until the 4:1 molar ratio was achieved. The *Mamu*-A*001 Tat_28-35_SL8 monomer was tetramerized to streptavidin-BV605 (0.1mg/mL, BD biosciences) or streptavidin-APC (0.25mg/mL, Agilent Technologies) at 8:1 monomer:streptavidin molar ratios for both fluorochromes. The tetramerization protocol was identical to that described above.

### In vitro characterization of NK cells

Using freshly isolated PBMC isolated from whole blood samples, NK cells were characterized using the antibodies, staining panel, and methods described in table 2 of our previous manuscript (20). Flow cytometric analysis was performed on a BD Symphony A3 (Becton Dickinson, Franklin Lakes, NJ), and the data were analyzed using FlowJo software for Macintosh (version 10.7.1).

### In vitro characterization of SIV-specific CD8 T cell phenotype and function

For all characterization of CD8 T cell phenotypes and functional analyses performed in this study, frozen PBMC isolated from whole blood samples were used. The panels used are indicated in Tables II-IV.

For all panels, tetramer staining was performed prior to additional surface and intracellular staining. Tetramer stains were performed at room temperature in the dark for 45 minutes in RPMI media supplemented with 10% FBS and 50nM dasatinib (Thermo Fisher Scientific, Waltham, MA). After 45 minutes, the cells were washed in a solution of FACS buffer (2% FBS in a 1X PBS solution) containing 50nM dasatinib and surface stains were performed using the antibodies indicated in Tables II-IV for 20 minutes at room temperature in the dark. Cells were fixed in a 2% paraformaldehyde solution. Following a 20-minute incubation, samples were either run on a BD Symphony A3, or permeabilized and stained for 20 minutes at room temperature in medium B (Thermo Fisher Scientific, Waltham, MA) for intracellular markers. Flow cytometric analysis was performed as described above.

### Intracellular cytokine staining (ICS) assay

Intracellular cytokine staining (ICS) assays were performed as previously described, with slight modifications for optimal responses (20, 36). Briefly, frozen PBMC were thawed and rested for approximately 4-6 hours in RPMI media containing 15% FBS. Then, Gag_181-189_CM9 or Tat_28-35_SL8 peptides, or the SIVmac239 Gag peptide pool were added to the cells at a final concentration of 0.5 μg/mL. Cells and peptides were incubated at 37°C for 90 minutes, then 1 μg/mL Brefeldin A (BioLegend, San Diego, CA) and 2 μM monensin (BioLegend, San Diego, CA) were added along with CD107a-BV605, and incubated for 16 hours (overnight) at 37°C and 5% CO2. The following day, the cells were stained according to methods described above with the surface and intracellular antibodies described in Table IV. Flow cytometric analysis was performed as described above.

#### Statistical analysis

For statistical analyses performed with the same individuals across time, repeated measures ANOVA non-parametric tests were performed, with Dunnett’s multiple comparisons. For individuals with missing samples for a subset of timepoints, mixed-effects ANOVA tests were performed using geisser-greenhouse correction.

For statistical analysis comparing two groups of animals from the same timepoint, Mann-Whitney tests were used.

## Results

### Treatment with N-803 did not suppress chronic SIV replication in vaccinated or unvaccinated rhesus macaques

Our initial goal was to test the hypothesis that prophylactic vaccination elicits CD8 T cells that are recalled during N-803 treatment and suppress plasma viremia. We designed a study with 15 rhesus macaques expressing the *Mamu*-A1*001 MHC class I allele, but did not express the *Mamu*-B*008 or -B*017 alleles associated with SIV viral control (27, 38). Ten animals were vaccinated with a DNA prime/rAd5 boost regimen to generate a high frequency of SIV-specific CD8 T cells (Fig. 1, red and light blue). We co-administered a plasmid expressing rhesus macaque IL-12 along with the DNA vaccine in five of the animals (Fig. 1, light blue). All vaccine vectors expressed the SIVmac239 *gag* gene. Five animals remained unvaccinated (Figure 1, dark blue). This specific vaccine strategy was chosen to elicit CD8 T cells, but not prevent infection with SIV (32, 39). All 15 animals were then infected intravenously with 10,000 infectious units (IU) of the molecularly barcoded SIVmac239M strain (Figure 1) (34).

**Fig. 1.**
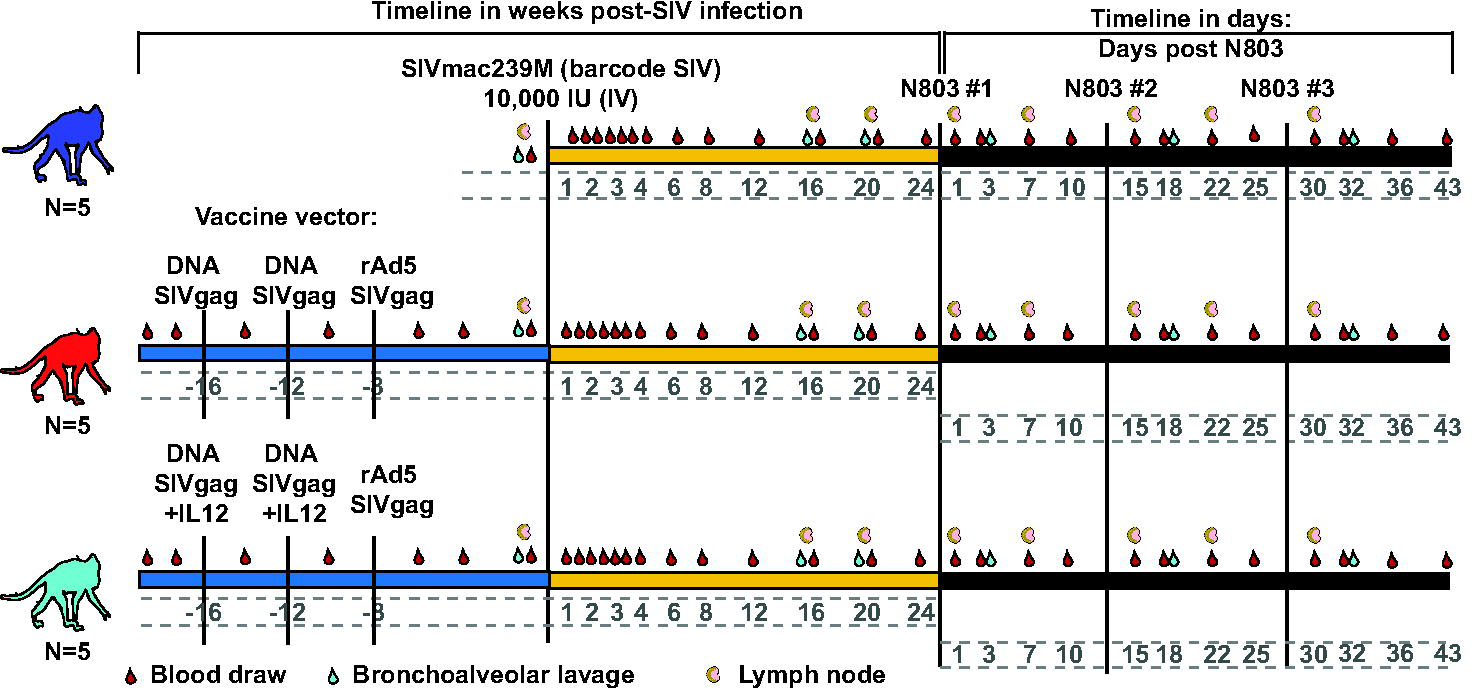
Outline for A*01+ macaque vaccine study. A*01+ rhesus macaques were either left unvaccinated (blue), or vaccinated with a heterologous prime/boost regimen with (light blue) or without (red) an IL-12 DNA vector adjuvant as indicated. Animals were infected with SIVmac239M for ∼6 months, then treated with 3 doses of 0.1mg/kg N-803, delivered subcutaneously, separated by two weeks per dose. Samples were collected as indicated.

As expected, the vaccine did not protect from SIV infection. However, the viral load set point in three of the 15 animals was below 10^4^ copies/mL (Fig. 2A). These three animals were all vaccinated. In contrast, chronic plasma viremia in the other 12 animals was between 10^5^ and 10^7^ copies/mL (Fig. 2A). Approximately 6 months after infection, all 15 animals received 3 doses of 0.1mg/kg N-803 separated by 14 days each (Fig. 1). N-803 treatment did not affect the plasma viral loads for any of the animals (Fig. 2A).

**Fig. 2.**
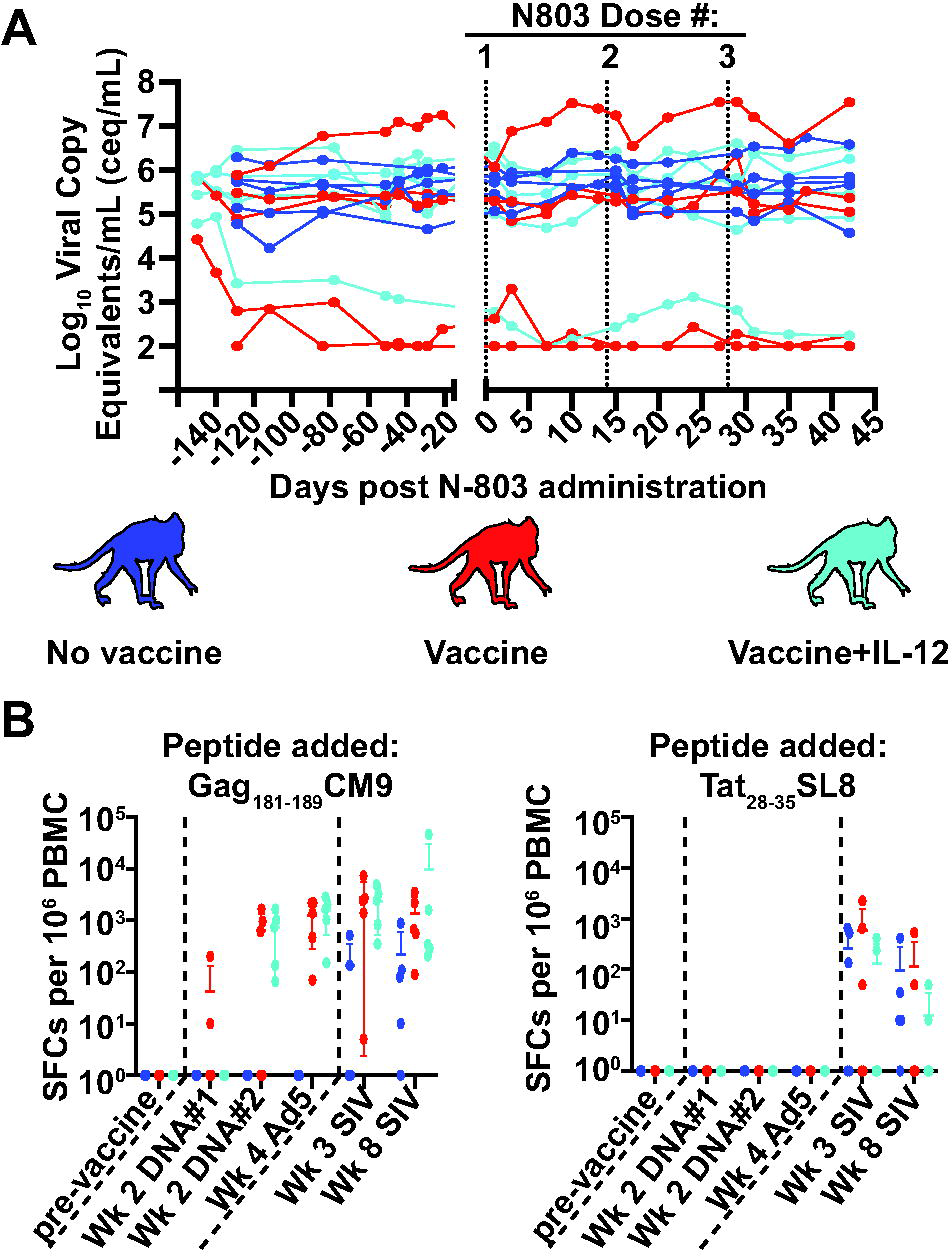
Viral loads for vaccinated and unvaccinated macaques are unchanged and do not differ during N-803 treatment. A, Plasma was isolated from whole blood samples from the unvaccinated (blue) and vaccinated (red and light blue) animals from the indicated timepoints post N-803 administration, and the Log_10_ virus copy equivalents/mL (ceq/mL) were determined as described in the methods. Vertical dashed lines indicate a timepoint on which N-803 was delivered. B, IFNγ ELISpots were performed on frozen PBMCs as indicated in the methods. Briefly, PBMC from the indicated timepoints post-vaccination or -SIV infection were thawed, then incubated overnight with 1 μM of the indicated peptides, or media alone as a negative control. Plates were developed according to the manufacturer’s instructions. A positive response was defined as the number of spot-forming colonies (SFCs) per 10^6^ PBMCs that were 2 standard deviations above the average value for the negative control, or 50 SFCs/10^6^ PBMC, whichever was greater.

We performed IFNγ-ELISpot assays using PBMC collected pre- and post-vaccination, as well as pre- and post-SIV infection (Fig. 2B) to confirm that the vaccine elicited Gag-specific T cells. The majority of vaccinated animals had a positive IFNγ ELISpot response when stimulated with the Gag_181-189_CM9 peptide by two weeks after the second DNA vaccination, and all animals had detectable Gag_181-189_CM9-specific T cells at 4 weeks after vaccination with the rAd5 particles (Fig. 2B, left graph). The unvaccinated macaques only produced Gag_181-189_CM9-specific T cells after SIV infection (Fig. 2B, left graph). No animals had detectable T cells specific for Tat_28-35_SL8 until after SIV infection (Fig. 2B, right graph).

### Vaccination did not affect the frequency of SIV-specific cells in lymph nodes after N-803 treatment

One attractive feature of N-803 and other IL-15 agonist complexes as an immunotherapeutic agent for HIV/SIV+ individuals is their ability to increase the trafficking of CD8 T cells to the lymph nodes (21, 22, 40). We performed flow cytometry on PBMC and lymph node samples to measure the absolute number (PBMC, Fig. 3A) and frequency (lymph nodes, Fig. 3B) of SIV-specific CD8 T cells. We included *Mamu*-A*001 tetramers presenting Gag_181-189_CM9 or Tat_28-35_SL8 peptides. The Gag_181-189_CM9 tetramer identifies cells that were initially elicited by vaccination, while the Tat_28-35_SL8 tetramer identifies cells that were elicited solely during SIV infection. The gating schematic for determining the frequencies of tetramer positive and negative cells is shown in supplementary figure 1A.

**Fig. 3.**
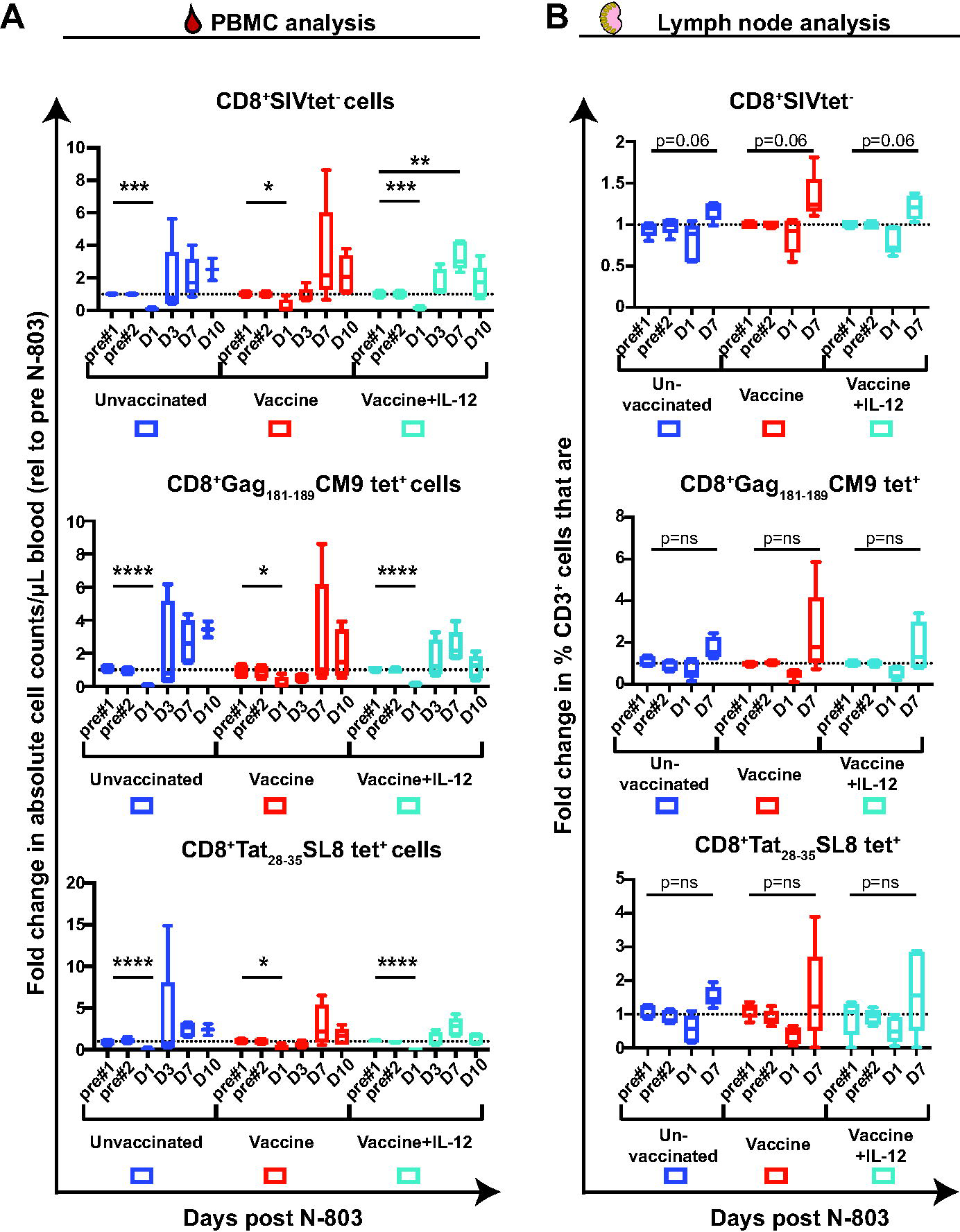
Vaccination does not change N-803 mediated trafficking of CD8 T cells. A, Frozen PBMC that were isolated from whole blood samples collected from the indicated timepoints pre- and post-N-803 treatment were stained for flow cytometric analysis as indicated in Table II. Complete white blood cell counts (CBC) were used to quantify the absolute number of CD3+ cells that were CD8+tetramer-(top), Gag_181-189_CM9 tetramer^+^(middle), and Tat_28-35_SL8 tetramer^+^ (bottom) cells per μL of blood. The results were then normalized to the average value for the absolute counts for each cell population from pre-N-803 controls and are displayed as the fold change in absolute cell counts/μL blood. Repeated measures ANOVA non-parametric tests were performed, with Dunnett’s multiple comparisons for individuals across multiple timepoints. For individuals for which samples from timepoints were missing, mixed-effects ANOVA tests were *p≤0.005; **p≤0.005;***, p≤0.0005; ****, p≤0.0001. B, Lymph nodes were collected from the indicated timepoints pre- and post-N-803 treatment. Cells were stained from frozen samples as indicated in (A) for flow cytometric analysis. The frequencies of CD3+MR1tetramer-T cells that were CD8+SIV tetramer- (top panel), CD8+Gag_181-189_CM9 tetramer^+^(middle), or CD8+Tat_28-35_SL8 tetramer^+^ (bottom) were determined, then the data were normalized to the average of the pre N-803 frequencies. Statistical analysis was performed as described in (A). p=ns; not significant.

**Table II.** Cytolytic granule/proliferation panel

| Antibody | Clone | Tetramer/Surface/Intracellular |
| --- | --- | --- |
| Gag<sub>181-189</sub>CM9-PE or Nef<sub>137-146</sub>RL10-PE | --- | Tetramer |
| Tat<sub>28-35</sub>SL8-BV605 | --- | Tetramer |
| MR1 5OPRU-BV421 | --- | Tetramer*** |
| CD3-AF700 | SP34-2 | Surface |
| CD4-BUV737 | SK3 | Surface |
| CD8-BUV395 | RPA-T8 | Surface |
| CD28-BV510 | CD28.2 | Surface |
| CD95-PE Cy7 | DX2 | Surface |
| CCR7-BV711 | G043H7 | Surface |
| LIVE/DEAD Near IR ARD | --- | --- |
| ki-67-AF488 | B56 | Intracellular |
| GranzymeB-PE CF594 | GB11 | Intracellular |
| Perforin-AF647 | PF344 | Intracellular |

\*\*\*The MR1 5OPRU tetramer staining was used to characterize MAIT cells, which were excluded from this manuscript.

Consistent with other studies, we found a decline in CD8 T cells in the peripheral blood one day after treatment with N-803 (21, 22). This decline was observed among bulk, Gag_181-189_CM9 tet+, and Tat_28-35_SL8 tet+ CD8 T cells (Fig. 3A) regardless of vaccination. However, we did not observe any statistically significant changes in the frequency of SIV-specific cells in the lymph nodes on days 1 or 7 post N-803 treatment (Figure 3B, top, middle, and bottom panels) for any of the animals. There was a trend towards increased bulk CD8 T cells in the lymph nodes, but this was not significant.

### N-803 alters the frequency of CXCR3+, but not CXCR5+, CD8 T cells in the peripheral blood in a vaccination-independent manner

We quantified the frequency of bulk and virus-specific CD8 T cells expressing the chemokine receptor CXCR5, CXCR3, and CCR6 during N-803 treatment. CXCR5 is associated with lymph node homing and localization (41, 42), and CXCR3 and CCR6 have been shown to mediate homing to sites of immune activation in the tissues (43, 44). Gating schematics for all chemokine markers can be found in supplementary fig. 2.

We found that CXCR5 was expressed on 1-20% of all CD8 T cells prior to N-803 treatment (Fig. 4A). This varied across animals and by antigen specificity, so we normalized the frequency of CXCR5+ cells at a specific time point to the pre-treatment average frequency for each individual animal that was plotted in Figure 4A. We did not observe any statistically significant increases in the frequency of CXCR5+ cells for the SIV tetramer-negative (Fig. 4B, top panel), Gag_181-189_CM9 tet+ (Fig. 4B, middle panel) or Tat_28-35_SL8 tet+ (Fig. 4B, bottom panel) cells on days 1, 3, 7, or 10. Consistent with the findings of Webb and colleagues (22, 22), we also found no statistically significant changes in the frequency of CXCR5+ cells in the lymph nodes (data not shown).

**Fig. 4.**
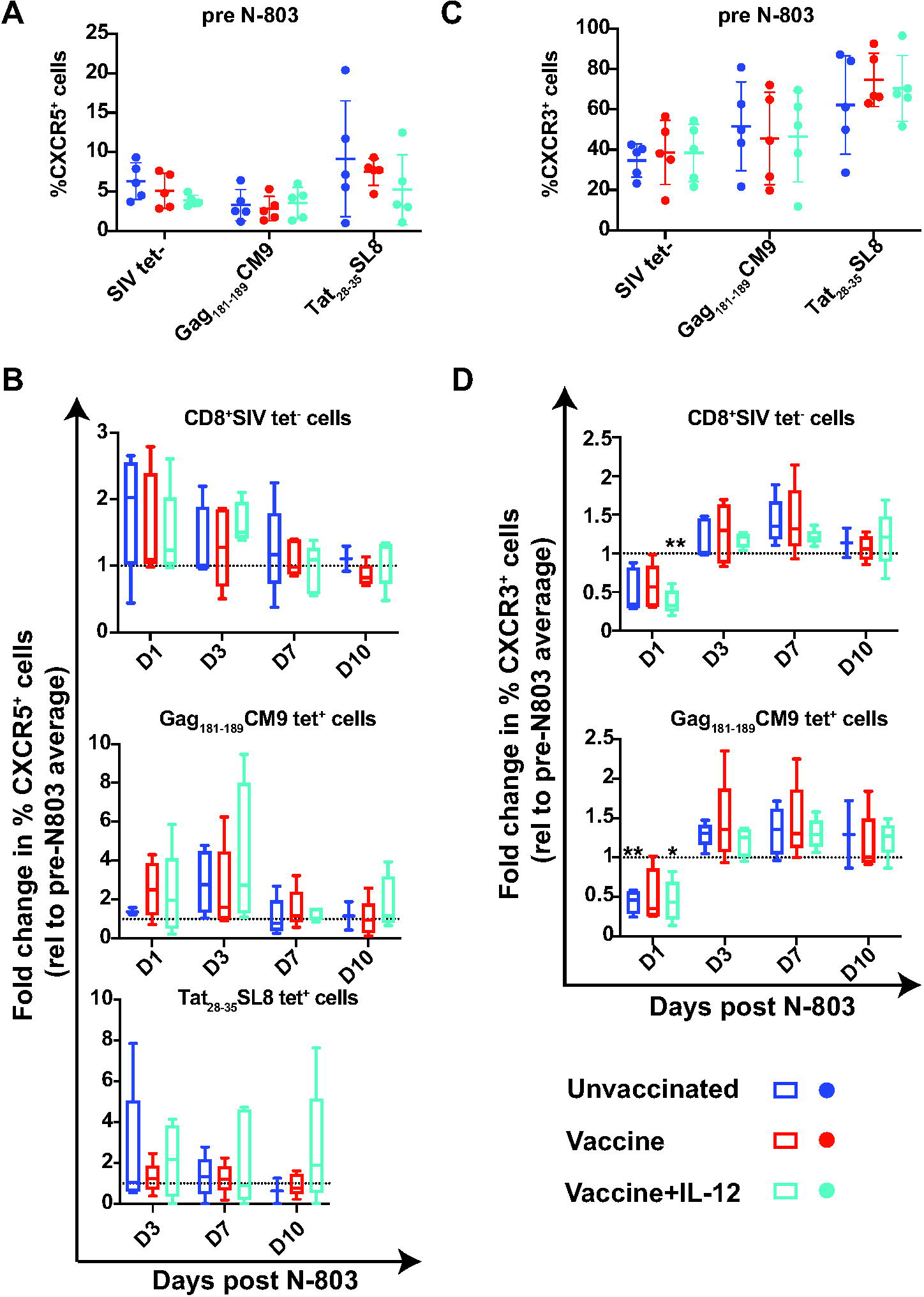
N-803 treatment does not change the frequency of CXCR5+ CD8 T cells, but does lead to a rapid decline in the frequency of CD8+CXCR3+ cells. A, Frozen PBMC were stained with the tetramers and antibodies indicated in Table III, and flow cytometry was performed, as described in the methods. Shown are the frequencies of CXCR5+ cells prior to N-803 treatment for CD8+ cells that were SIV tetramer-, Gag_181-189_CM9 tetramer^+^, or Tat_28-35_SL8 tetramer^+^ for each cohort. B, The fold change in the frequency of CXCR5+ cells relative to the pre N-803 average for CD8+SIV tetramer- (top), Gag_181-189_CM9 tetramer^+^ (middle), or Tat_28-35_SL8 tetramer^+^ (bottom) cells were determined for each timepoint post N-803 treatment. Repeated measures ANOVA non-parametric tests were performed, with Dunnett’s multiple comparisons for individuals across multiple timepoints. For individuals for which samples from timepoints were missing, mixed-effects ANOVA tests were performed using Geisser-Greenhouse correction. C, Frozen PBMC were stained as described in (A). Shown are the frequencies of CXCR3+ cells prior to N-803 treatment for CD8+ cells that were SIV tetramer-, Tat_28-35_SL8 tetramer^+,^ or Gag_181-189_CM9 tetramer^+^. D, The fold change in the frequency of CXCR3+ cells relative to the pre N-803 average for CD8+SIV tetramer- (top), or Gag_181-189_CM9 tetramer^+^ (bottom) cells were determined for each timepoint post N-803 treatment. Statistical analysis was performed as 0.05; **, p≤0.005.

**Table III.** Chemokine/trafficking panel

| Antibody | Clone | Tetramer/Surface/Intracellular |
| --- | --- | --- |
| Gag<sub>181-189</sub>CM9-PE | --- | Tetramer |
| Tat<sub>28-35</sub>SL8-BV605 | --- | Tetramer |
| MR1 5OPRU-APC | --- | Tetramer*** |
| CD3-AF700 | SP34-2 | Surface |
| CD4-BUV737 | SK3 | Surface |
| CD8-BUV395 | RPA-T8 | Surface |
| CXCR3-BV650 | G025H7 | Surface |
| CCR6-PE Cy7 | 11A9 | Surface |
| CXCR5-PerCP Efluor710 | MU5UBEE | Surface |
| CD122-BV421 | Mik-β3 | Surface |
| CD132-BB515 | AG184 | Surface |
| LIVE/DEAD Near IR ARD | --- | --- |

\*\*\*The MR1 5OPRU tetramer staining was used to characterize MAIT cells, which were excluded from this manuscript.

**Table IV.**
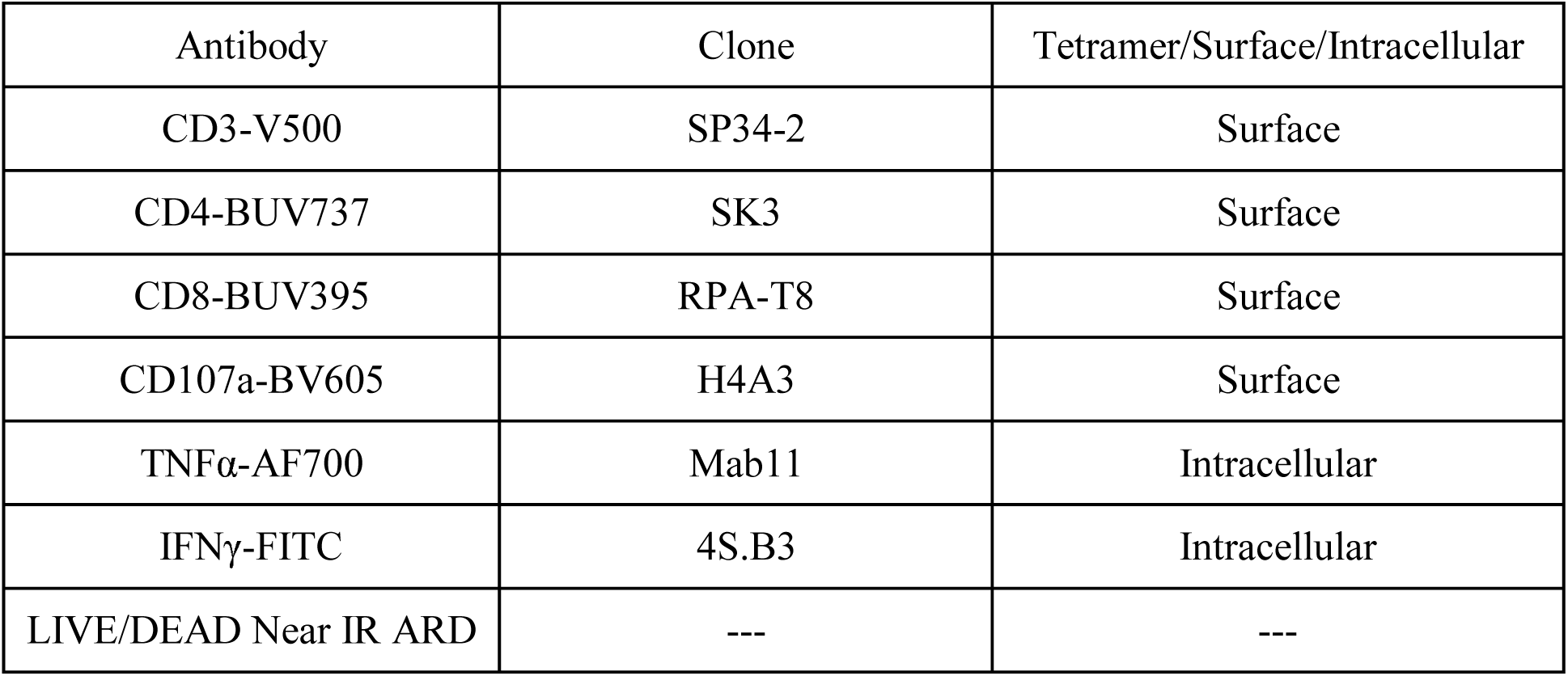
Antibodies used for ICS assay.

| Antibody | Clone | Tetramer/Surface/Intracellular |
| --- | --- | --- |
| CD3-V500 | SP34-2 | Surface |
| CD4-BUV737 | SK3 | Surface |
| CD8-BUV395 | RPA-T8 | Surface |
| CD107a-BV605 | H4A3 | Surface |
| TNFα-AF700 | Mab11 | Intracellular |
| IFNγ-FITC | 4S.B3 | Intracellular |
| LIVE/DEAD Near IR ARD | --- | --- |

We found that the frequency of CD8+ T cells expressing CXCR3 varied even more widely than those expressing CXCR5 (20-80%) prior to N-803 treatment (Fig. 4C). Interestingly, we observed a rapid decrease in the frequency of CXCR3+CD8+tetramer- and Gag_181-189_CM9 tet+ cells in the peripheral blood one day post-N-803, resulting in up to a 50% loss in the frequency of CXCR3+ cells relative to the pre-N-803 frequency (Fig. 4D, top and bottom panels). This decline was statistically significant relative to pre-N-803 timepoints for unvaccinated macaques (blue bars, Fig. 4D) and the macaques vaccinated with IL-12 plasmid (light blue bars), but not for the vaccinated animals who did not receive IL-12 plasmid (red bars). We did not analyze the frequency of CXCR3+ cells for the Tat_28-35_SL8 tet+ parent population due to very low numbers of these cells present in the peripheral blood on day 1 post N-803 (data not shown).

### Re-evaluation of N-803 efficacy based on prior immune mediated viral control

Prior vaccination did not predict whether macaques would respond to N-803 treatment, suggesting that this was not the sole factor responsible for the N-803-mediated virus suppression we observed in our previous study (Ellis-Connell et al., 2018, #55998). All the animals in our previous study spontaneously controlled SIV earlier during infection, a result typically associated with cytotoxic T cell function (45). Therefore, we decided to restructure the current study to test the alternative hypothesis that prior spontaneous SIV control predicts whether N-803 treatment improves CD8 T cell function with the potential to control SIV replication. Along the same lines, we expected that CD8 T cells from SIV non-controllers would not exhibit improved function after N-803 treatment.

To test this alternative hypothesis, we rearranged our animal groups into SIV non-controllers and controllers based on the viral load set point established prior to treatment with N-803. We included the 12 SIV non-controllers from this study (Fig. 5A, purple) whose viral load setpoint was above 10^4^ copies/ml. There were only 3 animals with a viral load set point lower than 10^4^ copies/ml in the current study, so we included samples from the 4 animals who responded to N-803 from the 2018 study to increase the size of the controller group (Ellis-Connell et al., 2018, #55998) (Fig. 5A, gold). We only included PBMC collected during the first 7 days after the first dose of N-803 treatment from the 2018 study (Fig. 5B) because the two studies differed after this time point.

**Fig. 5.**
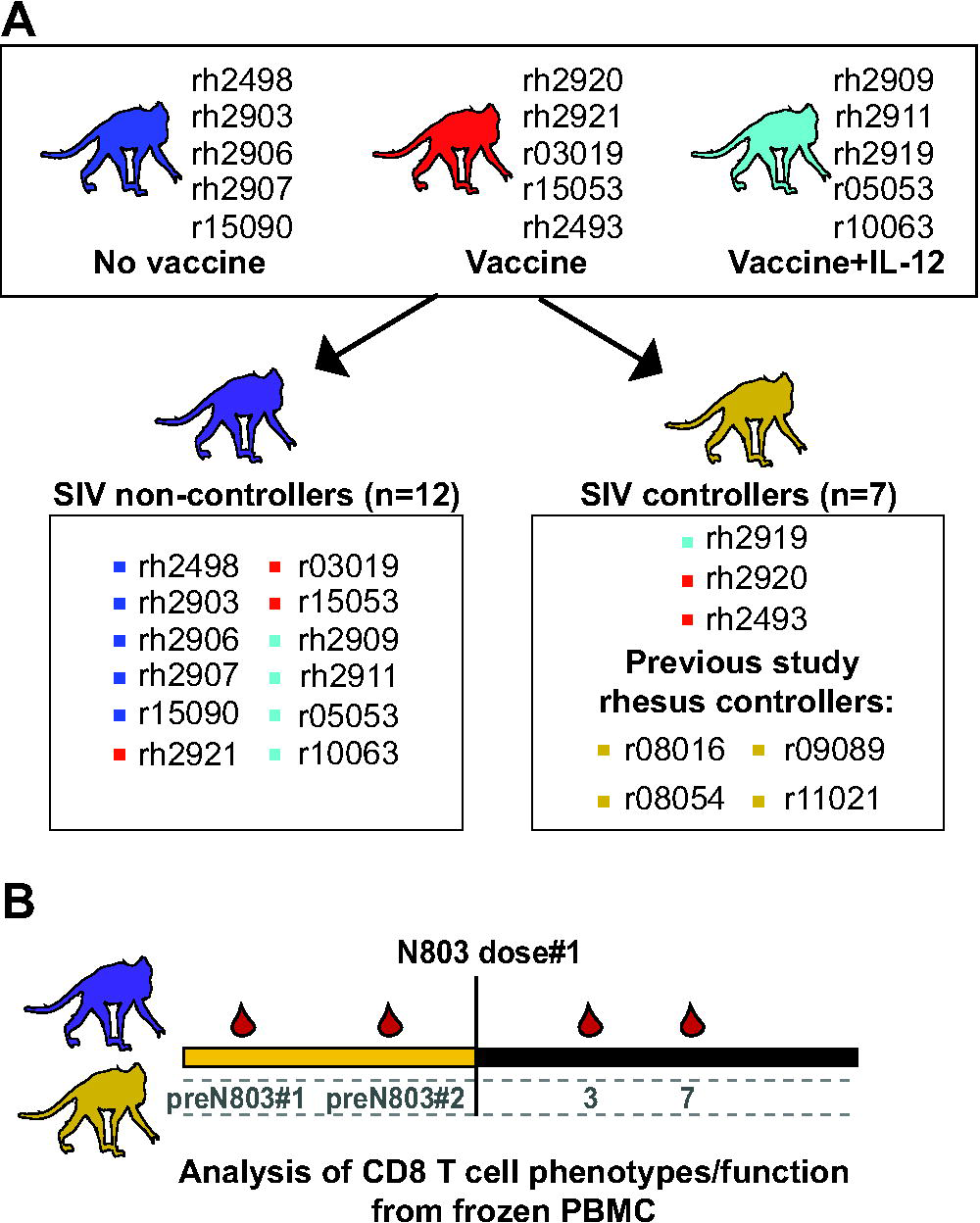
Rearrangement of animals into SIV controllers and SIV non-controllers. A, Animals from the original vaccine study (blue, red, and light blue) were grouped according to viral loads. SIV non-controllers (n=12, purple monkey) are shown in the box on the left and had viral loads above 10e4 ceq/mL (shown in Fig. 2). SIV controllers (n=7, gold monkey) are shown in the box on the right and had viral loads at or below 10e4 ceq/mL ((20); Fig. 2). B, Timeline of samples used for studies comparing CD8 T cells from SIV controllers and non-controllers. Frozen PBMC collected from the timepoints indicated from the animal groups described in (A) are shown on the timeline, and were used for downstream analysis.

### SIV controller status does not impact N-803 mediated increases in bulk CD8 T cells or NK cells

We measured the increases in absolute CD8 T cells (Fig. 6A) and NK cells (Fig. 6C) in the peripheral blood from SIV controllers and non-controllers. There was no difference in the N-803 mediated increase in either cell type between controllers (gold) and non-controllers (purple) 7 days after the first dose of N-803. We found similar results when we extended this analysis to the same cell populations for all three doses of N-803 used in the animal study described here (Figs. 6B, 6D).

**Fig. 6.**
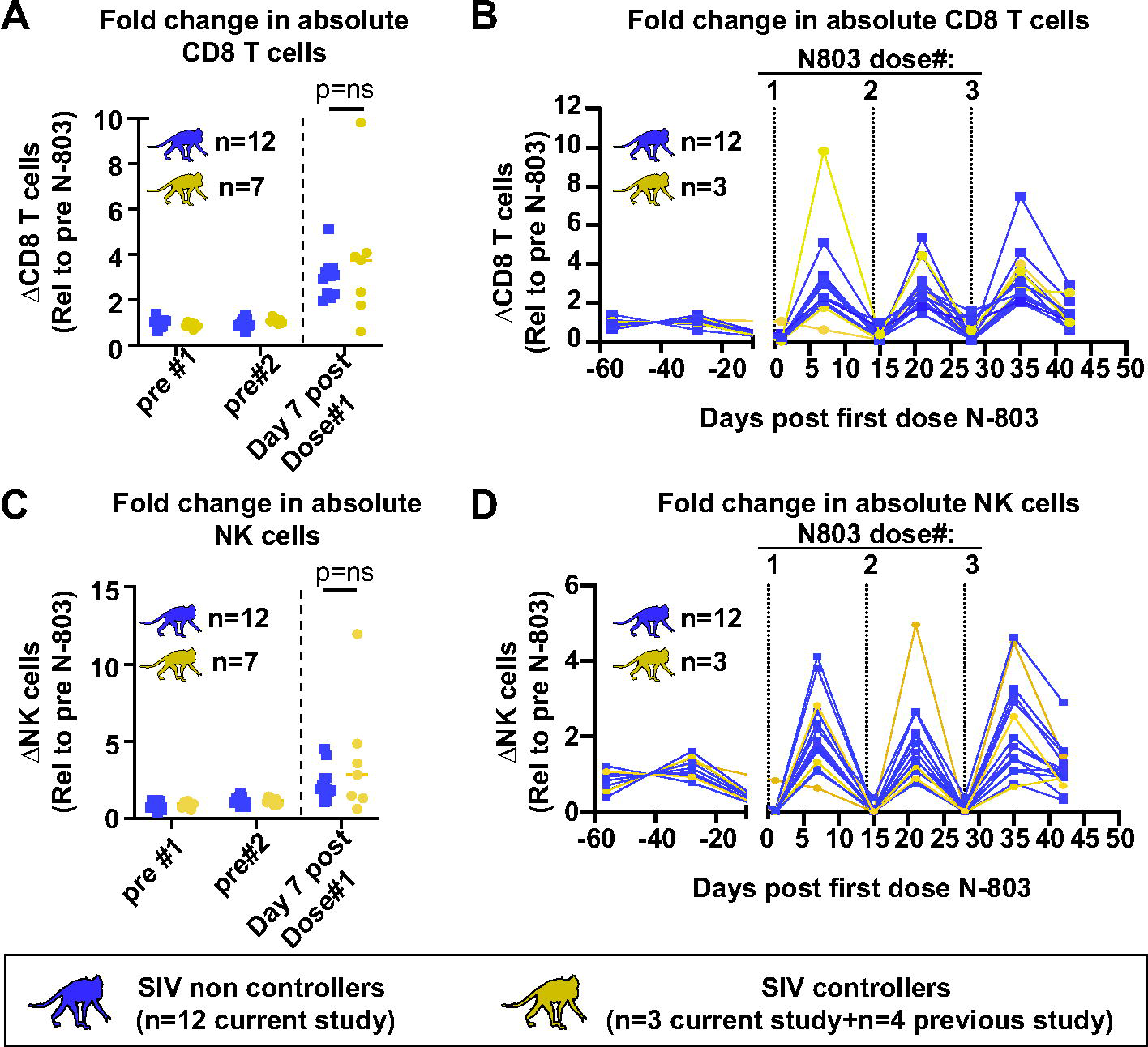
No differences in increases of bulk CD8 T cells or NK cells between SIV controllers and non-controllers during N-803 treatment. A-D. PBMC from SIV non-controllers (purple) or SIV controllers (gold) collected from the indicated timepoints post N-803 were stained with the panel originally described in Table III of (20). The frequency of CD8 T cells (defined as CD3+CD8+ cells; A and B) and NK cells (defined as CD3-CD8+NKG2A+ cells; C and D) were determined. Complete white blood cell counts (CBC) were used to quantify the absolute number of each cell population. Then, the data were normalized to the average value for the pre-treatment controls for each population, and are displayed as the fold change in absolute cell counts relative to the pre-treatment average. For figures A) and C), Mann-Wittney tests were performed to determine statistical significance. p=ns; not significant.

### SIV controller status does not impact N-803 mediated expansion of SIV-specific cells or alter their memory phenotypes

We wanted to determine if the virus-specific CD8 T cells from the SIV controllers were uniquely more responsive to treatment with N-803, when compared to their non-controller counterparts. To test this, we again utilized the *Mamu*-A*001 tetramers Gag_181-189_CM9 and Tat_28-35_SL8. We also included the *Mamu*-B*008-tetramer Nef_137-146_RL10 for the animals who expressed *Mamu*- B*008, but not *Mamu*-A*001 (r08016, r09089, and r08084 from Fig 5). See supplementary figure 1 for gating schematics.

Within each group, we examined the frequencies of CD3+ cells that were CD8+SIV tetramer negative (CD8+SIVtet-), Gag_181-189_CM9/Nef_137-146_RL10 tetramer positive, or Tat_28-35_SL8 tetramer positive (Fig. 7A). There were somewhat higher frequencies of CD3+CD8+SIV tet negative T cells in the SIV non-controllers compared to the SIV controllers at each timepoint and their frequencies were unaffected by N-803 treatment (Fig. 7A, left graph). However, when we examined the SIV tetramer+ cells, there were no statistically significant differences between the two groups in the frequencies of any tetramer+ cells before or after N-803 treatment (Fig. 7A, middle and right graphs). We also calculated the absolute number of Gag_181-189_CM9, Nef_137-146_RL10, and Tat_28-35_SL8+ CD8 T cells pre- and post-N-803 treatment (Fig 7B). While N-803 did increase the absolute number of these antigen-specific T cells in the peripheral blood 7 days after receiving N-803, the fold change in antigen-specific T cells was similar for both SIV controllers and non-controllers (Fig 7C). Although *in vivo* IL-15 treatment has been shown to preferentially expand central and effector memory cells (22, 46), we found no remarkable differences in frequencies of memory cells between the two cohorts of animals prior to N-803, nor did the memory phenotypes for each parent population of cells change significantly after N-803 treatment (data not shown).

**Fig. 7.**
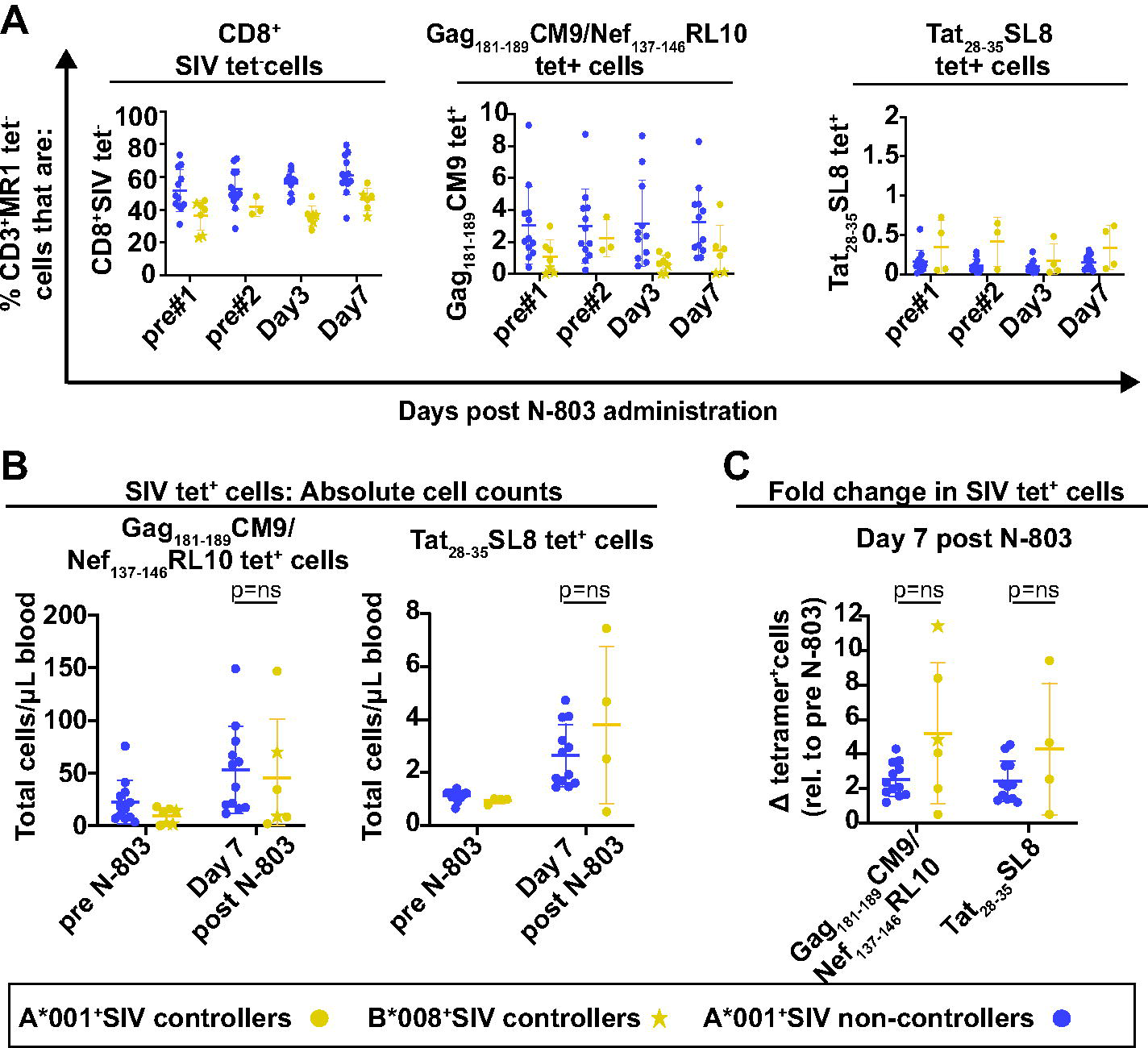
Frequencies of SIV-specific cells or memory populations between SIV controllers and non-controllers do not differ, and are not altered during N-803 treatment. A, Frozen PBMC from SIV non-controllers (purple) and SIV controllers (gold) that were collected from the indicated timepoints post N-803 were thawed, then stained with the panel described in Table III of the methods. Flow cytometric analysis was performed to determine the frequencies of CD3+MR1 tetramer-cells that were: CD8+SIV tet-(left panel), CD8+Gag_181-189_CM9 (gold/purple circles) or CD8+Nef_137-146_RL10 (gold stars) tet+ (middle panel); or CD8+Tat_28-35_SL8 tet+ (right panel) for each timepoint. B, Frozen PBMC from SIV non-controllers (purple) or SIV controllers (gold) collected from the indicated timepoints as described above. Complete white blood cell counts (CBC) were used to quantify the absolute number of each cell population. Then, the data were normalized to the average value for the pre-treatment controls for each population, and are displayed as the fold change in absolute cell counts relative to the pre-treatment average. Mann-Whitney tests were performed to determine statistical significance. p=ns, not significant.

### The frequencies of vaccine-elicited T cells expressing granzyme B and Ki-67 are increased more in SIV controllers compared to non-controllers

We hypothesized that the vaccine-elicited CD8 T cells from SIV controllers may exhibit improved function after N-803 treatment, when compared to non-controllers. This includes increased proliferation and cytolytic potential. Proliferation of cells can be measured by examining the frequency expressing ki-67, which is an intracellular marker that aids in cell division (47). N-803 is known to increase the frequency of ki-67+ CD8 T cells from HIV-naïve humans *in vitro* and SIV+ macaques *in vivo* (18, 20, 21), Granzyme B and perforin are molecules involved in degranulation and destruction of infected target cells (48, 49). N-803 and other IL-15 agonists increase the expression of granzyme B and perforin in NK cells and CD8 T cells in healthy individuals (18, 50, 51). We do not know how chronic immune activation, a feature common among SIV non-controllers (52), reduces the expansion of CD8 T cells producing Ki67, granzyme B, or perforin upon N-803 treatment, when compared to SIV controllers.

We measured the frequency of CD8+SIV tetramer+ and CD8+tetramer-negative cells expressing granzyme B and ki-67 before and after receiving N-803 (gating is shown in supplementary fig 3). We found the frequency of Gag_181-189_CM9 tetramer+ EM and TM cells producing ki-67 or granzyme B was increased most notable on day 3 after N-803 treatment (Figs. 8A and 9A, left two graphs). The fold increases in Gag_181-189_CM9 tetramer+ cells expressing these markers was most apparent in the SIV controllers, compared to the SIV non-controllers (Figs. 8A and 9A, right two graphs). This was attributed to the fact that the baseline frequencies of Gag_181-189_CM9 tetramer+ EM and TM cells expressing ki-67 or granzyme B was much higher in the SIV non-controllers when compared to controllers (Figs 8 and 9A, left panels, and Supplementary Fig. 3). This is likely a result of ongoing antigenic stimulation by circulating virus. Central memory cells were too rare to characterize (data not shown).

**Fig. 8.**
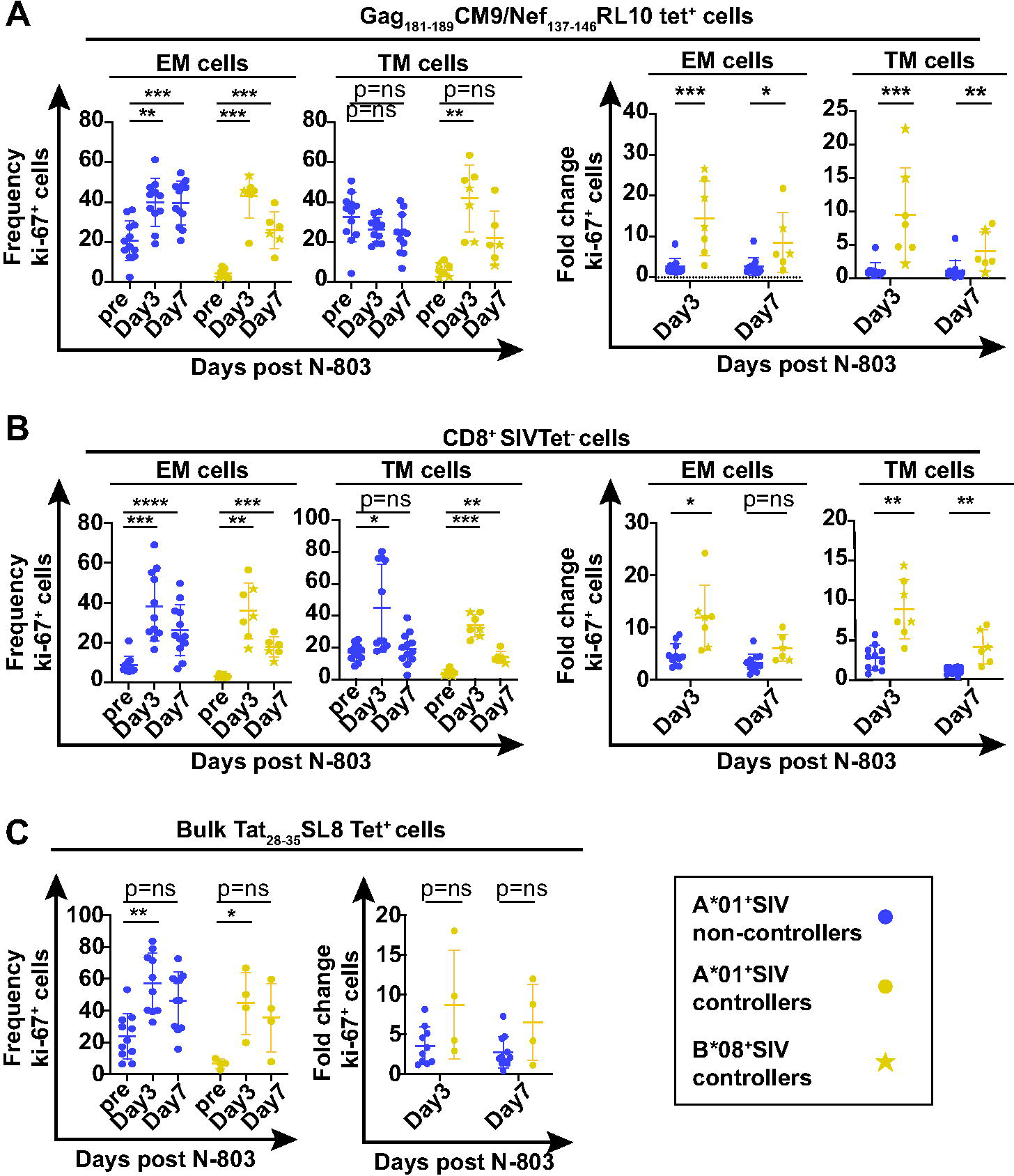
N-803 treatment increases the frequency of SIV-specific cells expressing the proliferation marker ki-67 to a greater extent in SIV controllers compared to SIV non-controllers. Frozen PBMC from SIV non-controllers (purple) and SIV controllers (gold circles and stars) that were collected from the indicated timepoints post N-803 were thawed, and flow cytometry was performed using the panel described in Table II of the methods. The frequencies of ki-67+ cells (left panels) or the fold change in ki-67 relative to pre N-803 controls (right panels) were examined for the indicated time points on effector memory (CD28-CD95+CCR7-; EM), and transitional memory (CD28+CD95+CCR7-; TM), cells for the following parent populations of cells: A) CD8+Gag_181-189_CM9 (gold/purple circles) or CD8+Nef_137-146_RL10 (gold stars) tet+ cells or B) CD8+SIV tet-cells. C, CD8+Tat_28-35_SL8 tet+ cells were analyzed in bulk for the frequency (left panel) and fold change (right panel) of ki-67+ cells. For graphs of ki-67 frequencies within a cohort, repeated measures ANOVA non-parametric tests were performed, with Dunnett’s multiple comparisons for individuals across multiple timepoints. For graphs comparing the fold change in percent ki67+ cells between controllers and non-controllers for a given time point, Mann-Whitney tests were performed to determine statistical significance: p=ns, not significant *, p≤0.05; **, p≤0.005 ***, p≤0.0005.

**Fig. 9.**
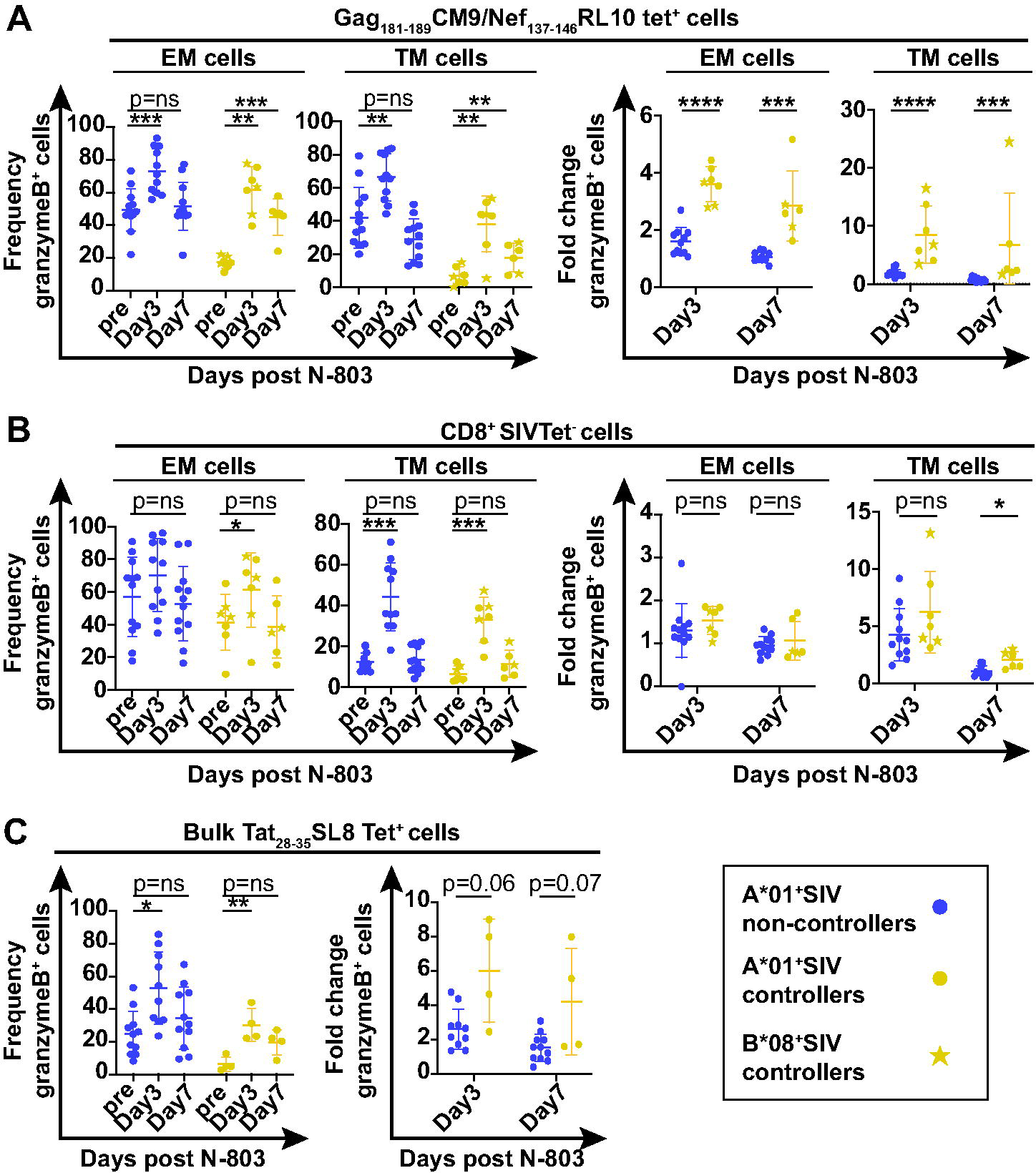
N-803 treatment increases the frequency of SIV-specific cells expressing the proliferation marker granzyme B to a greater extent in SIV controllers compared to SIV non-controllers. Frozen PBMC from SIV non-controllers (purple) and SIV controllers (gold circles and stars) that were collected from the indicated timepoints post N-803 were thawed, and flow cytometry was performed using the panel described in Table II of the methods. The frequencies of granzyme B+ cells (left panels) or the fold change in granzyme B+ cells relative to pre N-803 controls (right panels) were examined for the indicated time points on effector memory (CD28-CD95+CCR7-; EM), and transitional memory (CD28+CD95+CCR7-; TM), cells for the following parent populations of cells: A) CD8+Gag_181-189_CM9 (gold/purple circles) or CD8+Nef_137-146_RL10 (gold stars) tet+ cells or B) CD8+SIV tet-cells. C, CD8+Tat_28-35_SL8 tet+ cells were analyzed in bulk for the frequency (left panel) and fold change (right panel) of granzyme B+ cells. For graphs of granzyme B frequencies within a cohort, repeated measures ANOVA non-parametric tests were performed, with Dunnett’s multiple comparisons for individuals across multiple timepoints. For graphs comparing the fold change in percent granzyme B+ cells between controllers and non-controllers for a given time point, Mann-Whitney tests were performed to determine statistical p=ns, not significant; *, p≤0.05; **, p≤0.005; ***, p 0.0005.

In contrast to the Gag_181-189_CM9 tetramer+ cells, the changes in the frequencies of CD8+ SIV tetramer negative EM and TM cells expressing ki-67 or granzyme B after N-803 treatment were similar in all animals (Figs. 8B and 9B, left two graphs). This resulted in fewer differences between SIV non-controllers and SIV controllers. The population of Tat_28-35_SL8 tetramer+ cells was analyzed in bulk as there were too few cells to examine individual memory populations. The frequency of Tat_28-35_SL8 tetramer+ cells expressing ki-67 and granzyme B increased similarly after N-803 treatment across animals (Figs. 8C and 9C).

### N-803 treatment of SIV controllers increases the frequency of virus-specific CD8 T cells expressing CD107a

We performed intracellular cytokine staining (ICS) assays using PBMC collected pre- and post-N-803 treatment. We compared the frequency of antigen-specific CD8 T cells producing TNFα, IFNγ, and CD107a between SIV controllers and non-controllers. A representative gating schematic for CD107a, TNFα, and IFNγ is shown in supplementary figure 4 for an SIV controller (gold, top panels) and an SIV non-controller (purple, bottom panels).

We found that all animals had an increased frequency of CD8+CD107a+, IFNγ+, and TNF+ cells in response to Gag or Nef peptides, or a Gag peptide pool, relative to unstimulated controls (Supplementary Fig. 4B). We normalized the data from stimulated to unstimulated controls collected from the matched timepoints (Fig. 10A-C).

**Fig. 10.**
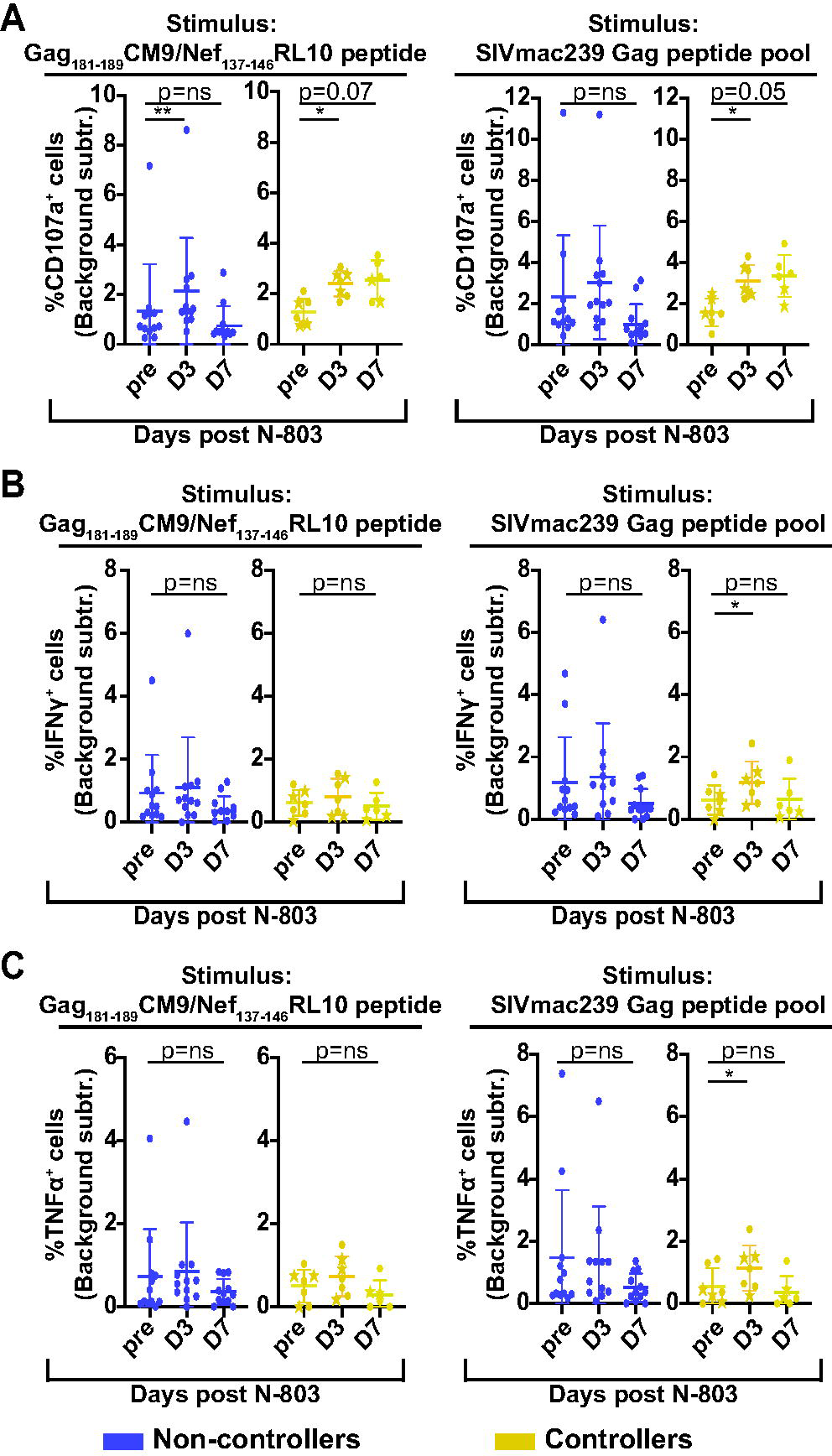
N-803 treatment in SIV controllers, but not SIV non-controllers, leads to an improvement in CD107a production in functional assays. Frozen PBMC from SIV non-controllers (purple) and SIV controllers (Gold) that were collected from pre N-803 (pre), Day 3 post N-803 (D3), and Day 7 post N-803 (D7) were thawed and incubated overnight with either media alone, 0.5ug/mL Gag_181-189_CM9, or Nef_137-146_RL10 peptides, or 0.5ug/mL of SIVmac239 Gag peptide pool. The next day, flow cytometry was performed using the panel described in Table IV of the methods. The data were normalized by background subtraction. The frequency of cells producing CD107a (A), IFNγ (B), or TNFα(C) after background subtraction in response to the indicated antigen are shown. For all statistical analyses, repeated measures ANOVA non-parametric tests were performed, with Dunnett’s multiple comparisons. For individuals for which samples from timepoints were missing, mixed-effects ANOVA tests were performed using Geisser-Greenhouse correction. *, p≤0.05; **, p≤0.005.

Similar to frequencies of CD8 T cells expressing ki-67 or granzyme B, the SIV non-controllers (purple) had a higher background of CD8 T cells expressing CD107a compared to controllers (Supplementary Fig. 4B). After eliminating the background signal, we found that N-803 treatment had little impact on the antigen-stimulated CD107a production from SIV non-controllers (Fig. 10A, purple). Similarly, after elimination of background signal, there were no statistically significant differences in antigen-stimulated IFNγ or TNFα production after N-803 treatment for the SIV non-controllers (Fig. 10B and C, purple).

In contrast, cells collected from SIV controllers at day 3 post-N-803 treatment had a statistically significant increase in the frequency of antigen-specific CD107a+ CD8 T cells. The frequency of CD107a+ antigen-specific cells collected from day 7 post-N-803 remained elevated, but not statistically significant (Fig. 10A, right panel, gold). We found minor but statistically significant increases in IFNγ+ (Fig 10B, right panel, gold) and TNFα+ (Fig 10C, right panel, gold) CD8 T cells with Gag peptide pool stimulation from cells collected from day 3 post-N-803 treatment compared to the pre-N-803 timepoint.

## Discussion

We previously found that administration of the IL-15 superagonist, N-803, to SIV+ ART-naïve vaccinated macaques led to rapid, but transient, control of plasma viremia (20). Here, we tested the hypothesis that vaccination, prior to SIV infection, was necessary for N-803 treatment to induce suppression of plasma viremia. Unfortunately, we found that prior vaccination, alone, did not confer N-803-mediated control of plasma viremia during N-803 treatment in macaques (Fig. 2). We observed that peripheral SIV-specific CD8 T cells from vaccinated and unvaccinated macaques exhibited similar changes in frequencies and phenotypes after N-803 treatment (Figs. 3-4). Furthermore, SIV-specific CD8 T cells from vaccinated animals did not exhibit improved targeting to the lymph nodes during N-803 treatment when compared to unvaccinated controls (Fig 3). These observations implied that prophylactic vaccination prior to SIV infection, by itself, did not produce a higher frequency of N-803 responsive CD8 T cells.

Another phenotype of previous N-803 responsive animals was that they spontaneously controlled SIV replication during the earlier stages of infection (53), and three of them expressed the *Mamu-B*008* MHC class I allele associated with viral control. Therefore, we wanted to test the hypothesis that a predisposition to viral control is associated with an improved response to N-803 treatment. We divided the animals into two groups based on their ability to spontaneously control early SIV infection. We found that N-803 treatment led to larger increases in the frequencies of ki-67 and granzyme B+ Gag_181-189_CM9 tetramer+ cells of SIV controllers compared to SIV non-controllers (Figs 8 and 9). Additionally, in SIV controllers but not non-controllers, CD8 T cells collected after N-803 treatment were more likely to produce CD107a after Gag peptide stimulation. These findings suggest that Gag-specific CD8 T cells from SIV controllers may have the potential to exhibit improved proliferation and cytotoxicity with N-803 treatment when compared to SIV non-controllers.

Our study could have important implications for the clinical use of N-803 to boost CD8 T cell function in HIV+ individuals. Those with uncontrolled plasma viremia have dysregulated HIV/SIV specific CD8 T cells (8, 52). We observed that the SIV-specific CD8 T cells of non-controllers had a chronically activated phenotype, with higher frequencies of granzyme B and ki-67+ cells before N-803 treatment (Figs. 8-9, Supplementary Fig. 3). N-803 did not lead to a substantial number of CD8 T cells with this activated phenotype during treatment (Figs. 8 and 9). Chronic immune activation leads to a type of immune “exhaustion” that leaves HIV/SIV specific cells unable to combat infection (52, 54), which is difficult to reverse through immunotherapeutic interventions alone. While antiretroviral treatment (ART) does not fully restore the function of HIV/SIV-specific CD8 T cells (55), ART-mediated control of plasma viremia may improve CD8 T cell function sufficiently to allow for a more efficacious response to N-803. Our data suggests that this type of chronic immune activation also weakens the responsiveness to N-803.

While vaccination did not always lead to control of SIV infection in our study, the vaccine-elicited Gag_181-189_CM9 tetramer+ cells from the SIV controllers exhibited greater increases in frequencies of ki-67 and granzyme B+ cells during N-803 treatment compared to the non-controllers (Figs. 8 and 9A). Furthermore, Gag-specific CD8 T cells in ICS assays from SIV controllers displayed a greater ability to produce CD107a compared to Gag-specific CD8 T cells from SIV non-controllers (Fig. 10). This improvement in proliferative and cytotoxic capability could suggest that vaccination, combined with N-803 immunotherapy, could impact the SIV reservoir. Webb and colleagues did not find that N-803 reduced the SIV reservoir, but this was in unvaccinated macaques (22). However, the fact that we observed differences in the function of CD8 T cells in the peripheral blood in the SIV controllers during N-803 treatment when compared to non-controllers (Figs. 8-10), could mean that vaccination plus N-803 treatment could impact CD8 T cell cytotoxic function in the lymph nodes and tissues. When comparing vaccinated to unvaccinated macaques, we did not observe differences in plasma viremia or changes in the frequencies of CD8 T cells present in the lymph nodes (Fig. 3). Due to poor cell yields from the lymph nodes SIV controllers at critical timepoints, we were unable to compare cell frequencies, phenotypes, and function of vaccine-elicited CD8 T cells from lymph node biopsies from SIV controllers and non-controllers. Since the lymph nodes are critical sites of SIV replication, it will be interesting to focus on the dynamics of N-803-boosted, vaccine-elicited CD8 T cells from SIV controllers and non-controllers in future, larger animal studies

One interesting observation was that CD8 T cells expressing the chemokine marker CXCR3 were significantly reduced one day after N-803 treatment from the peripheral blood (Fig. 4). We do not think that this was necessarily a result of the downregulation of CXCR3 surface expression. Rather, we hypothesize that the CXCR3+ CD8 T cells may have migrated away from the peripheral blood to the tissues. CXCR3 is a chemokine typically involved in the trafficking of T cells to sites of immune activation, such as tissues (43). Studies involving other IL-15 agonists indicate that IL-15 induces migration of CD8 T cells to the tissues *in vivo,* and also explains how successful N-803 has been in the treatment of tissue-localized cancers (56–58).

CXCR3+ CD8 T cells also have enhanced cytotoxic function upon stimulation with IL-15 *ex vivo* (59). If N-803 causes an increase in the migration of highly cytotoxic SIV specific T cells to the tissues, this could have implications for the elimination of the HIV/SIV reservoir in tissues. Although one macaque study suggested that N-803 did not reduce the reservoir in lymph nodes in SIV+, ART-suppressed macaques (22), there may be other scenarios where CXCR3+ CD8 T cells can control or reduce the reservoir. For example, a recent study found that HIV+ individuals treated with N-803 as part of a clinical trial had a small but significant reduction in CD4 T cells with inducible virus in the peripheral blood after N-803 treatment (23). These individuals were also all on ART at the time of N-803 treatment. Neither of these studies focused on tissues such as the gut, a known reservoir for HIV/SIV (60, 61). Thus, future studies could examine if N-803 boosted CD8 T cells are capable of migrating to tissue sites of HIV/SIV infection and reducing the reservoir.

Overall, our study begins to dissect out the host conditions under which N-803 may be most effective at improving the cytotoxic function of CD8 T cells. We found that prophylactic vaccination alone did not condition animals to respond to N-803. However, animals predisposed to viral control had a population of CD8 T cells that were more responsive to N-803, when compared to animals that never controlled virus replication. Specifically, it appears as though N-803 increases the function of the SIV-specific CD8 T cells from SIV controllers to a greater degree than SIV non-controllers. Our findings support that N-803 could have great potential as an immunotherapeutic agent for HIV/SIV+ individuals, particularly in the setting of HIV/SIV control, such as during ART treatment.

## Acknowledgments

We thank the NIH Tetramer Core Facility (contract number 75N93020D00005) for providing the Mamu B*008 Nef_137-146_RL10 and Mamu A*001 Tat_28-35_SL8 biotinylated monomers. We thank Drs. John Mascola, Robert Seder, and Wing-Pui Kong for the hCMV/R-SIVmac239 *gag* DNA vaccine vector. We also thank Drs. George Pavlakis and Barbara Felber for the rhesus IL-12 plasmid vector. We finally thank Dr. Brandon Keele for the use of the SIVmac239M barcoded virus.

We thank staff at the Wisconsin National Primate Resource Center (WNPRC) for excellent veterinary care of the animals involved in this study.

## SUPPLEMENTARY FIGURE LEGENDS

**Supplementary Fig. 1.**
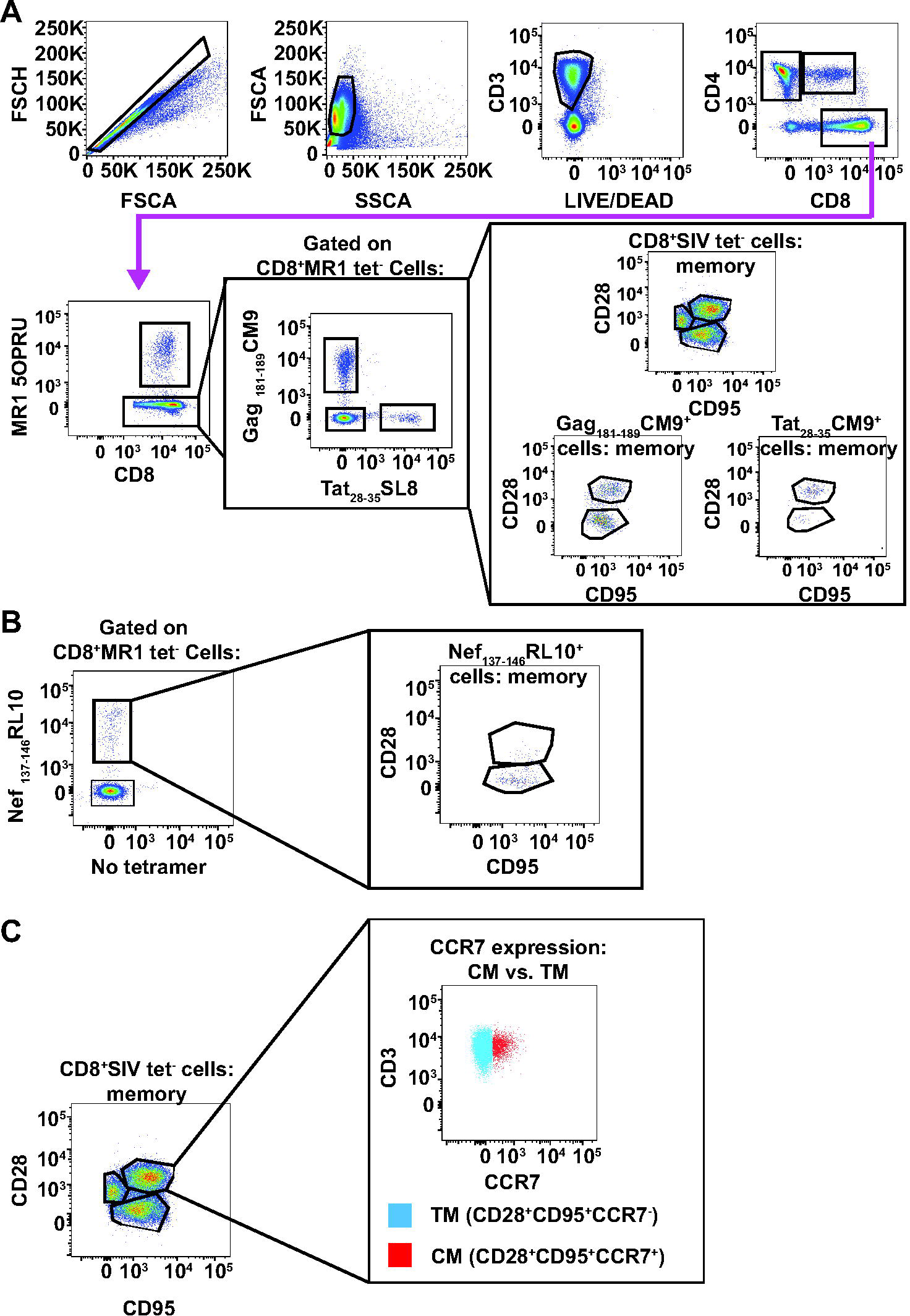
Representative gating schematic for Tetramer+ cells and memory markers in A*001+ and B*008+ macaques. A, Frozen PBMC from all A*001+ macaques in this study were stained with the antibodies indicated in Table II of the methods. Flow cytometry was performed as described in the methods. Shown is a representative gating schematic for CD8+Gag_181-189_CM9 tetramer+,CD8+Tat_28-35_SL8 tetramer+, and CD8+SIV tetramer-cells, and memory sub-populations for each respective tetramer-positive or -negative parent population. B, Representative gating schematic for Nef_137-146_RL10 tetramer+ cells in B*008+ macaques. Cells were stained identically to the samples in (A) except Nef_137-146_RL10 tetramer was used instead of Gag_181-189_CM9 tetramer, and no Tat_28-35_SL8 tetramer was used. Shown is a representative sample indicating Nef_137-146_RL10 tetramer+ cells and the memory subpopulations of those cells. C, Gating schematic used to define Effector memory, Central memory, and Transitional memory cells. Memory subpopulations that were shown in (A) and (B) were further differentiated into effector memory (CD28-CD95+CCR7-), transitional memory (CD28+CD95+CCR7-) or central memory (CD28+CD95+CCR7+) based on CCR7 expression. Central memory and transitional memory cell phenotypes were validated by examining CCR7 expression based on the CD28+CD95+ parent gate.

**Supplementary Fig. 2.**
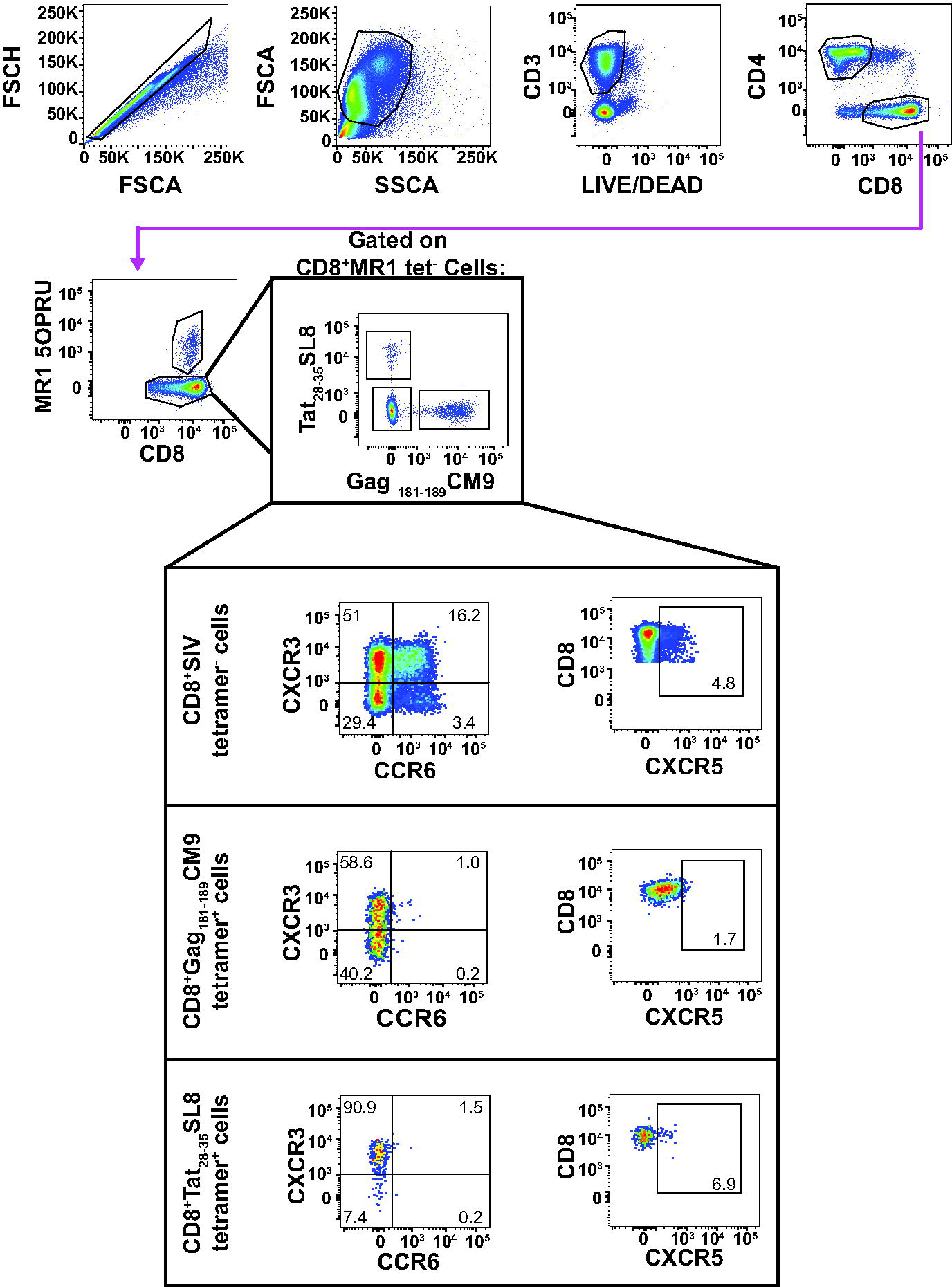
Representative gating schematic for CXCR3, CCR6, and CXCR5 expression on CD8 T cells. Frozen PBMC were stained with the antibodies indicated in Table III of the methods. Flow cytometry was performed as described in the methods. Shown is a representative gating schematic for CD8+Gag_181-189_CM9 tetramer+,CD8+Tat_28-35_SL8 tetramer+, and CD8+SIV tetramer-cells, and their respective CXCR3+, CCR6+, and CXCR5+ subpopulations.

**Supplementary Fig. 3.**
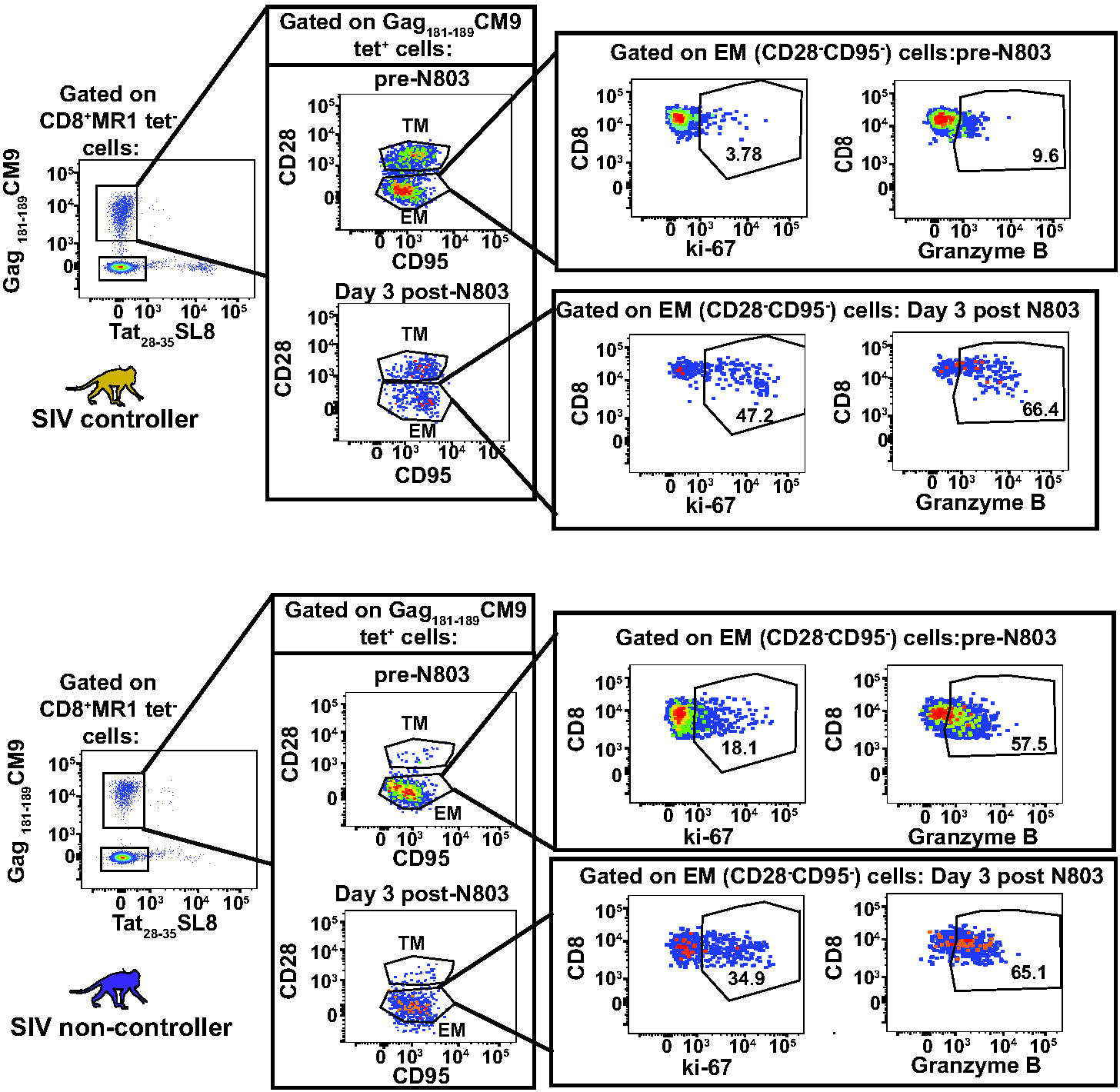
Representative gating schematic for ki-67 and Granzyme B expression on SIV specific cells of SIV controllers and non-controllers. Frozen PBMC from SIV controllers (gold; top panels) or SIV non-controllers (purple, bottom panels) were stained with the antibodies indicated in table II of the methods, and flow cytometry was performed. The frequency of ki-67+ and GranzymeB+ cells were determined for all memory subpopulations of CD8+SIV tetramer-, Gag_181-189_CM9 tetramer+, Nef_137-146_RL10 tetramer+, and Tat_28-35_SL8 tetramer+ cells. Shown is representative ki-67 and Granzyme B staining for the effector memory (EM) cells of Gag_181-189_CM9 tetramer+ cells of an SIV controller (gold) and an SIV non-controller (purple) from pre N-803 and Day 3 post N-803 timepoints.

**Supplementary Fig. 4.**
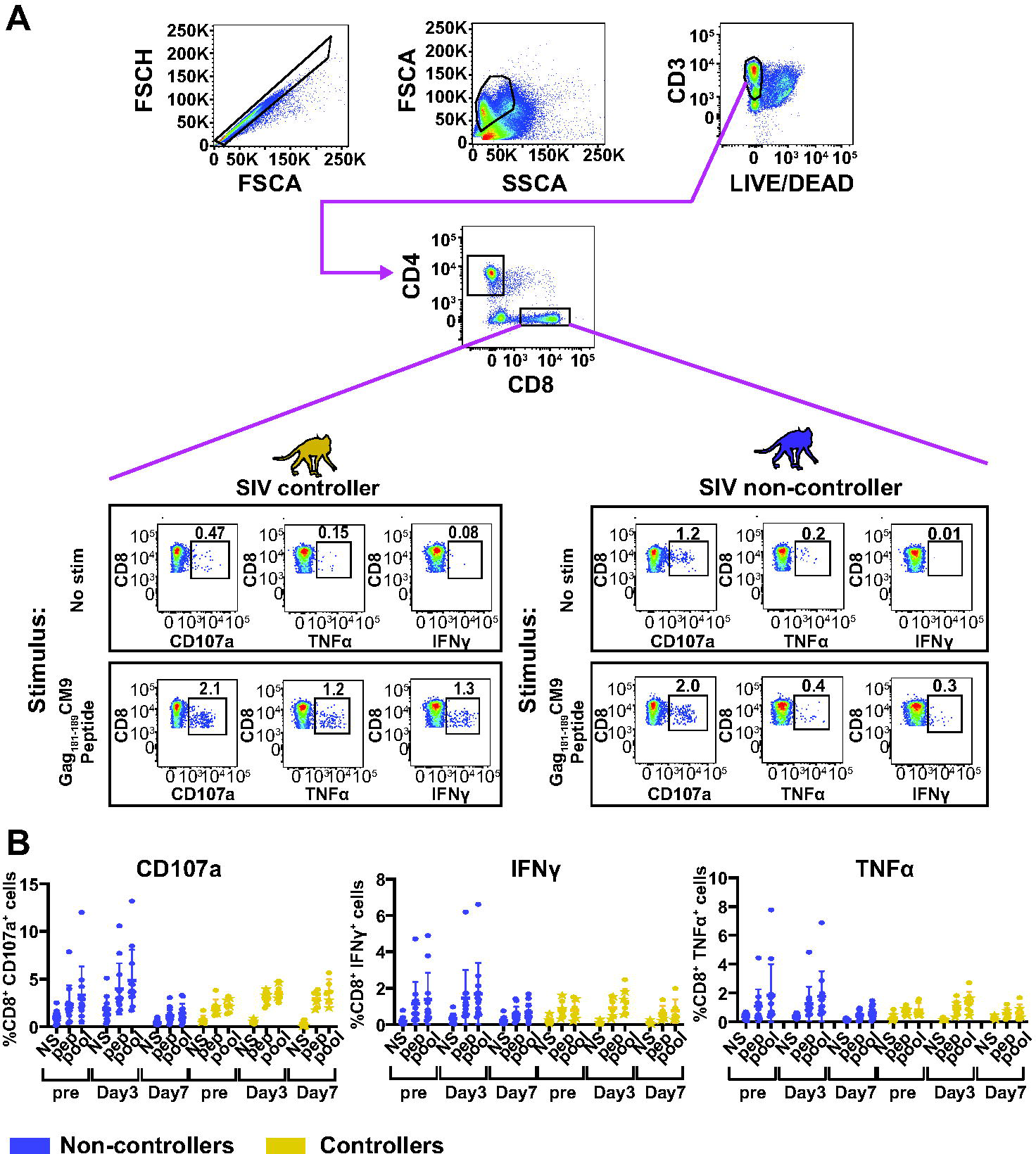
A, Representative gating schematic for intracellular cytokine staining (ICS) assay. Frozen PBMC from SIV non-controllers (purple) or SIV controllers (gold) were incubated overnight with media alone (no stim), or stimulated with peptides as indicated in the methods. For this representative figure, 0.5μg/mL of Gag_181-189_CM9 peptide was used. The following day, the cells were stained with the antibodies indicated in table IV of the methods, and flow cytometry was performed. Shown is a gating schematic for CD8+ T cells, and representative CD107a+, TNFα +, IFNγ + subpopulations with each respective stimulus for an SIV controller (gold) and an SIV non-controller (purple). B, Frequencies of CD107a+, IFNγ +, or TNFα + from intracellular cytokine staining assays from Fig. 10 prior to background subtraction. Intracellular cytokine staining assays were performed as indicated in Fig. 10 and cells were stained as indicated in table IV. Shown are the frequencies of CD8 T cells producing CD107a (left), IFNγ (middle), or TNFα (right) prior to background subtraction. NS; no stim, pep; μg/mL of Gag_181-189_CM9 or Nef_137-146_RL10 peptides, pool; 0.5ug/mL of SIVmac239 Gag peptide pool.

